# Myddosome clustering in IL-1 receptor signaling regulates the formation of an NF-kB activating signalosome

**DOI:** 10.1101/2023.01.06.522894

**Authors:** Fakun Cao, Rafael Deliz-Aguirre, Fenja H. U. Gerpott, Elke Ziska, Marcus J. Taylor

**Affiliations:** Max Planck Institute for Infection Biology, Chariteplatz 1, Berlin D-10117 Germany

## Abstract

Signaling pathways can produce digital invariant outputs and analog outputs that scale with the amount of stimulation. In IL-1 receptor (IL-1R) signaling both types of outputs require the Myddosome, a multi-protein complex. The Myddosome is required for polyubiquitin chain formation and NF-kB signaling. However, the ways in which these signals are spatially and temporally regulated to drive switch-like and proportional outcomes is not understood. We find that during IL-1R signaling, Myddosomes dynamically re-organize into large, multi-Myddosome clusters at the cell membrane. Blockade of Myddosome clustering using nanoscale extracellular barriers reduces NF-kB activation. We find that Myddosomes function as a scaffold that assembles an NF-kB signalosome consisting of E3-ubiquitin ligases TRAF6 and LUBAC, K63/M1-linked polyubiquitin chains, phospho-IKK, and phospho-p65. This signalosome preferentially assembles at regions of high Myddosome density, which enhances the recruitment of TRAF6 and LUBAC. Extracellular barriers that restrict Myddosome clustering perturbed the recruitment of both ligases. We found that LUBAC was especially sensitive to clustering with a sevenfold lower recruitment to single Myddosomes than clustered Myddosomes. This data reveals that the clustering behavior of Myddosome provides the basis for digital and analog IL-1R signaling.

## Introduction

Signaling pathways can give digital or ‘switch-like’ responses that are invariant (Shah and Sarkar 2011) or alternatively give analog responses that are proportional to the amount of stimulatory input (Nunns and Goentoro 2018). For instance in the innate immune system, IL-1 activation of NF-kB has both an invariant component and a response proportional to the stimulating dose (DeFelice et al. 2019; Son et al. 2021). Critical to these responses is the temporal and spatial control of reactants within a signaling pathway. Protein effectors must be brought together at a precise point within the cell to ensure accurate signal transduction. One way in which signaling pathways seem to achieve this is through the formation of signalosomes: subcellular compartments containing clusters of receptors and signaling effectors. Signalosomes are found in multiple receptor signaling systems such as tyrosine kinases, immune receptors and Wnt receptors (Case, Ditlev, and Rosen 2019). In the case of IL-1 signaling, an NF-kB signalosome assembles in response to stimulation (Tarantino et al. 2014). Thus, a key question is how signalosomes, such as that associated with NF-kB activation, can activate both invariant and proportional responses.

A critical component of many signalosomes are protein scaffolds that can bind and concentrate multiple signaling effectors (Jaqaman and Ditlev 2021; Wu 2013). The ability of protein scaffolds to oligomerize or self-assemble plays a crucial role in signalosome formation and downstream signaling (Ditlev, Case, and Rosen 2018). In the immune system, oligomeric protein scaffolds serve a central role in tuning the intensity and duration of signaling responses (Wu 2013). The Myddosome is an oligomeric complex that is crucial for IL-1R signal transduction and an inflammatory innate immune response. The Myddosome activates the generation of K63-ubiquitin linked (K63-Ub) chains via directly interacting with the E3 ligase TNF Receptor-Associated Factor 6 (TRAF6) (Ye et al. 2002). Myddosomes can also activate the generation of M1-Ubiquitin linked (M1-Ub) chains via the linear ubiquitin chain assembly complex (LUBAC) (Tokunaga et al. 2009). K63-Ub and M1-Ub chains recruit the IκB kinase (IKK) complex that activates NF-kB signaling and results in the translocation of the RelA NF-kB subunit to the nucleus (Wertz and Dixit 2010; Iwai 2012). This signaling pathway can encode both digital and analog outputs as defined by downstream readouts such as RelA dynamics or transcriptional responses (Tay et al. 2010; Hughey et al. 2015; Cheng et al. 2021). However, whether and how the Myddosome encodes both invariant and proportional outputs upstream of NF-kB, has not been investigated.

Here we address this problem using live cell imaging to visualize Myddosome formation and downstream signal transduction in response to IL-1 stimulation. We observe that Myddosomes reorganize into clusters or regions of the plasma membrane that contain a high density of complexes. Physically limiting Myddosome clustering with extracellular barriers diminishes NF-kB activation. We find that Myddosomes function as scaffolds that nucleate a signalosome containing K63/M1-Ub chains and markers of NF-kB activation. Single Myddosomes can nucleate the formation of this signalosome, suggesting it is an invariant or digital signaling output of the complex. However this NF-kB signalosome preferentially formed at clusters and the degree of Myddosome clustering proportionally increases the size of this signalosome. In particular clustering amplifies the production of M1-Ub. Live cell imaging revealed that the ubiquitin ligases TRAF6 and LUBAC are preferentially recruited to Myddosome clusters. Restricting clustering diminished TRAF6 recruitment and severely perturbed HOIL1 recruitment. We conclude that clustering is an important determinant of E3 ubiquitin ligase recruitment, and this dynamic encodes a signaling output that is proportional to the nanoscale density of complexes within the cluster. These results suggest a mechanism for how Myddosomes can encode both digital and analog responses upstream of NF-kB in IL-1R signaling.

## Results

### Myddosomes dynamically reorganize into clusters

Understanding how IL-1R and Myddosome encode digital and analog outputs requires understanding where these differences arise within the signaling network. Therefore to uncover the link between the spatial organization of Myddosomes and the production of downstream signaling outputs we used a supported lipid bilayer (SLB) system functionalized with IL-1 (Deliz-Aguirre et al. 2021) to visualize the dynamics of IL-1R-Myddosome signal transduction. We found that MyD88-GFP assembles into puncta at the cell surface (Fig. 1A). Initially MyD88-GFP puncta are spatially segregated, but over time MyD88-GFP puncta move and coalesce, forming brighter puncta, and eventually larger dense patch like structures at the cell-SLB interface (Fig. 1A, Movie 1). These dynamic clusters are similar to large Myddosome structures observed in Toll-like receptor 4 signaling that are associated with stronger NF-kB responses (Latty et al. 2018). We decided to investigate how these dynamic Myddosome clusters are involved in IL-1R signal transduction.

**Figure 1.**
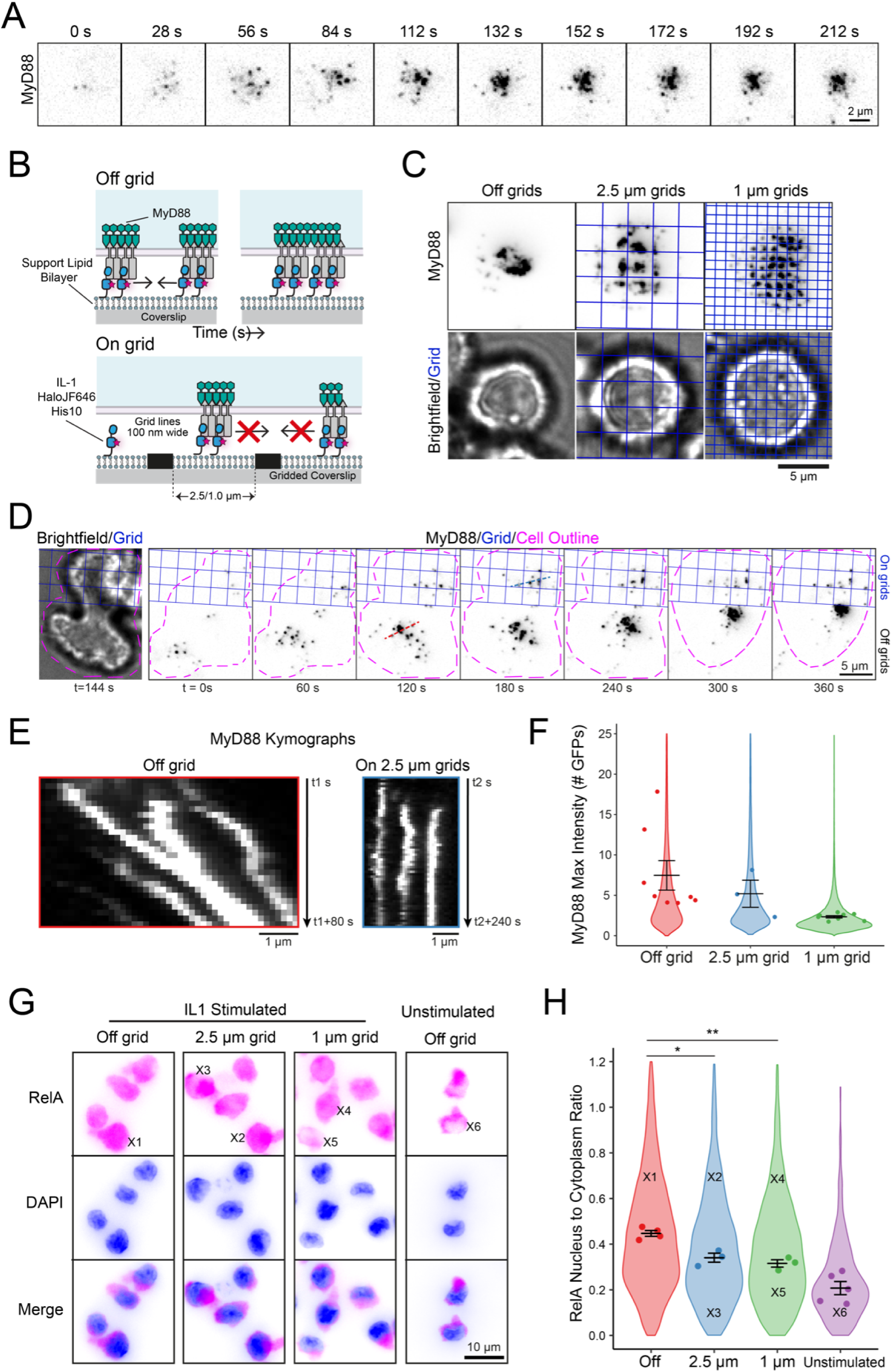
**Myddosomes are tethered to the cell surface and extracellular barriers inhibit Myddosome coalescence and diminish NF-kB activation.** (A) Timelapse TIRF microscopy images showing an EL4-MyD88-GFP cell interacting with a IL-1 functionalized SLB. MyD88-GFP assembles into puncta that over time cluster and coalesce at the cell:SLB interface. (B) Schematic of illustrating a work model for Myddosomes being tethered to the plasma membrane via interaction with IL-1R bound to IL-1. Based on this model, we predict physical barriers (on grid) would restrict the diffusion of IL-1 on the SLB and limit Myddosome clustering. With no external barriers present (off grid), MyD88 puncta can merge to form larger multi-complex assemblies. (C) TIRF and brightfield microscopy images of EL4-MyD88-GFP cells incubated for 30 mins with IL-1 functionalized SLBs formed off grid and on 1 and 2.5 µm grids. In the presence of a 1 and 2.5 µm grids, Myddosomes only coalesce within individual corrals and do not form multi-complex clusters. Scale bar, 5 µm. (D) Time series showing Myddosome formation in an EL4 cell interacting with both continuous and 2.5 µm gridded partitioned SLBs. Scale bar, 5 µm. (E) Kymographs from panel D showing the coalescence of MyD88-GFP puncta off grid and the restricted movement of MyD88-GFP puncta on 2.5 µm grids. Scale bar, 1 µm. (F) Quantification of MyD88-GFP puncta maximum fluorescence intensity normalized to GFP from cells stimulated off and on 2.5 or 1 µm grids, at a ligand density of 10 IL-1/µm^2^. Violin plots show the distribution of average max puncta intensities from individual cells across replicates. Data points superimposed on the violin plots are the averages from independent experimental replicates. The average max MyD88 puncta intensity (mean ± SEM): for off grid is 7.5 ± 1.8 GFPs, n = 8 replicates; for 2.5 µm grids is 5.2 ± 1.7 GFPs, n = 3 replicates; for 1 µm grids is 2.3 ± 0.1 GFPs, n = 8 replicates. 6-50 cells were measured per experimental replicate. Bars represent mean ± SEM. (G) Widefield images showing RelA localization in unstimulated EL4 cells and EL4 cells stimulated by SLB formed on and off grids. EL4 were fixed 30 min after addition to IL-1-functionalized SLBs and stained for RelA (magenta); DAPI stained nuclei (blue). Scale bar, 10 µm. (H) Quantification of RelA nucleus to cytoplasm ratio. Violin plots show the distribution of measurements from individual cells. Data points superimposed on the violin plots are the averages from independent experiments. The RelA nucleus to cytoplasm ratio of single cells marked with X in panel (G) are superimposed on the violin plot. RelA nucleus to cytoplasm ratio off grids, on 2.5 and 1 µm grids, and unstimulated conditions are 0.45 ± 0.01, 0.34 ± 0.02, 0.32 ± 0.02, 0.21 ± 0.03 (mean ± SEM) respectively. The p value are * = 0.0133 and ** = 0.0027. Bars represent mean ± SEM (n = 3 to 5 experimental replicates, with ≥89 cells quantified per replicate for each condition).

The dynamic clustering of MyD88 puncta within the plane of the cell membrane suggests that Myddosomes are tethered to the inner leaflet of the plasma membrane. This tethering is likely due to heterotypic TIR domain interactions between MyD88 and the IL-1R/IL-1RAcP complex (Nimma et al. 2017). We predict, if this model is correct, extracellular barriers that restrict the diffusion of SLB-tethered IL-1 would restrict Myddosome mobility and clustering (Fig. 1B). To test this model, we used coverslips nano-printed with chromium barriers arranged into multiple 0.5 mm square grids. Within these square grids, chromium grid lines were printed into 1 or 2.5 µm square corrals (Fig. S1A). The chromium grid lines function as physical barriers and create an array of corralled SLBs with uniform dimensions. We confirmed SLB formation within these grids and that IL-1 ligands are freely mobile within corrals, but diffusion between corrals is restricted (Fig. S1B). In cells which landed on nanopatterned grids, MyD88 puncta were confined to individual corrals (Fig. 1C). We imaged cells that straddled the boundary between the 2.5 µm grid and continuous coverslip, so we could analyze Myddosome dynamics on/off grids within the same cell (Fig. 1D, Movie 2). Kymograph analysis reveals that in the same cell only off grid MyD88 puncta clustered. In contrast, MyD88 puncta on the 2.5 µm grid were confined to individual corrals and did not merge with puncta in adjacent corrals (Fig. 1E). We conclude that Myddosomes are biochemically coupled to extracellular IL-1 via IL-1R, and extracellular barriers limit the diffusion of complexes within the plasma membrane.

To determine whether clustering regulates downstream signaling requires tools that can isolate a single Myddosome for comparative analysis to clustered Myddosomes. We analyzed the size distribution of MyD88 puncta on/off grids (FIg. 1F). Off grid MyD88 puncta had a broad size distribution and a mean MyD88 copy number of 7.5 ± 1.8 MyD88s (Fig. 1F), suggesting a mix of clusters and single complexes. However on 2.5 or 1 µm grids, the mean MyD88 copy number was 5.2 ± 1.7 or 2.3 ± 0.1 MyD88s (Mean ± SEM, Fig. 1F, also see methods). Only 9.3 ± 1.6% of MyD88 puncta on 1 µm grids had an intensity consistent with ≥1 Myddosome complexes and 1.9 ± 0.6% puncta had an intensity consistent with ≥2 Myddosome complexes (see methods and Fig. 1F). Thus the majority of MyD88-GFP puncta on 1 µm grids are small transient MyD88 assemblies or single Myddosome complexes (Deliz-Aguirre et al. 2021), and nano-patterned IL-1 functionalized SLBs can spatially isolate single Myddosomes. In conclusion, nano-patterned coverslips are an effective tool to assay how the spatial organization of Myddosomes is functionally connected to digital and analog signaling outputs.

### Inhibiting Myddosome clustering diminishes RelA translocation to the cell nucleus

We tested whether inhibition of clustering perturbed NF-kB signaling by measuring RelA translocation to the nucleus. In unstimulated cells (incubated with unfunctionalized SLBs), RelA staining is limited to the cytosol and depleted within the cell nucleus (Fig. 1G). In cells incubated with IL-1 functionalized SLBs without grids, RelA translocates from the cell cytosol to the nucleus, resulting in stronger nuclear staining (Fig. 1G). For cells on grids, a mixture of both events was observed (2.5 µm and 1 µm grids, Fig. 1G). When we quantify RelA translocation, we found that the nucleus to cytoplasm ratio of RelA staining significantly decreased on 1 or 2.5 µm grids (normalized RelA nucleus to cytoplasm ratio of 0.45 ± 0.01 off grid versus 0.34 ± 0.02 and 0.32 ± 0.02 on 2.5 and 1 µm grids respectively, mean ± SEM, Fig. 1H). We conclude that the inhibition of Myddosome clustering impacts NF-kB activation and RelA translocation to the nucleus. The implication of these results is that Myddosome dynamics and spatial density at the cell surface is linked to the production of signaling outputs required for NF-kB activation.

### Myddosomes colocalize with an NF-kB-activating signalosome composed of K63-Ub/M1-Ub poly-ubiquitin chains, phospho-IKK and phospho-p65

Innate immune signaling complexes are proposed to function as signaling scaffolds that recruit and activate downstream effectors (Wu 2013). We speculated that the spatial organization of a signaling complex could regulate its scaffolding function and this could be the basis for invariant or proportional signaling responses. This scaffolding model suggests spatial colocalization between Myddosomes and biochemical signaling reactions. We examined the colocalization of Myddoosmes with IL-1 signaling outputs such as K63-Ub, M1-Ub, phosphorylated IκB kinase (pIKK) complex and phosphorylated RelA subunit p65 (pp65) using immunofluorescence and TIRF microscopy (Fig. 2A). We found these signaling outputs had a punctate staining pattern that colocalized with dense patches of clustered MyD88-GFP puncta (Fig. 2A). Detailed analysis of these Myddosomes patches shows that MyD88 was organized into heterogenous puncta of different size and irregular shapes (Fig. 2B). While these puncta of K63-Ub, M1-Ub, pIKK and pp65 staining did not uniformly coat MyD88 patches, these structures were clearly associated with MyD88 clusters. These results confirm that downstream signaling outputs are generated at cell surface Myddosomes.

**Figure 2.**
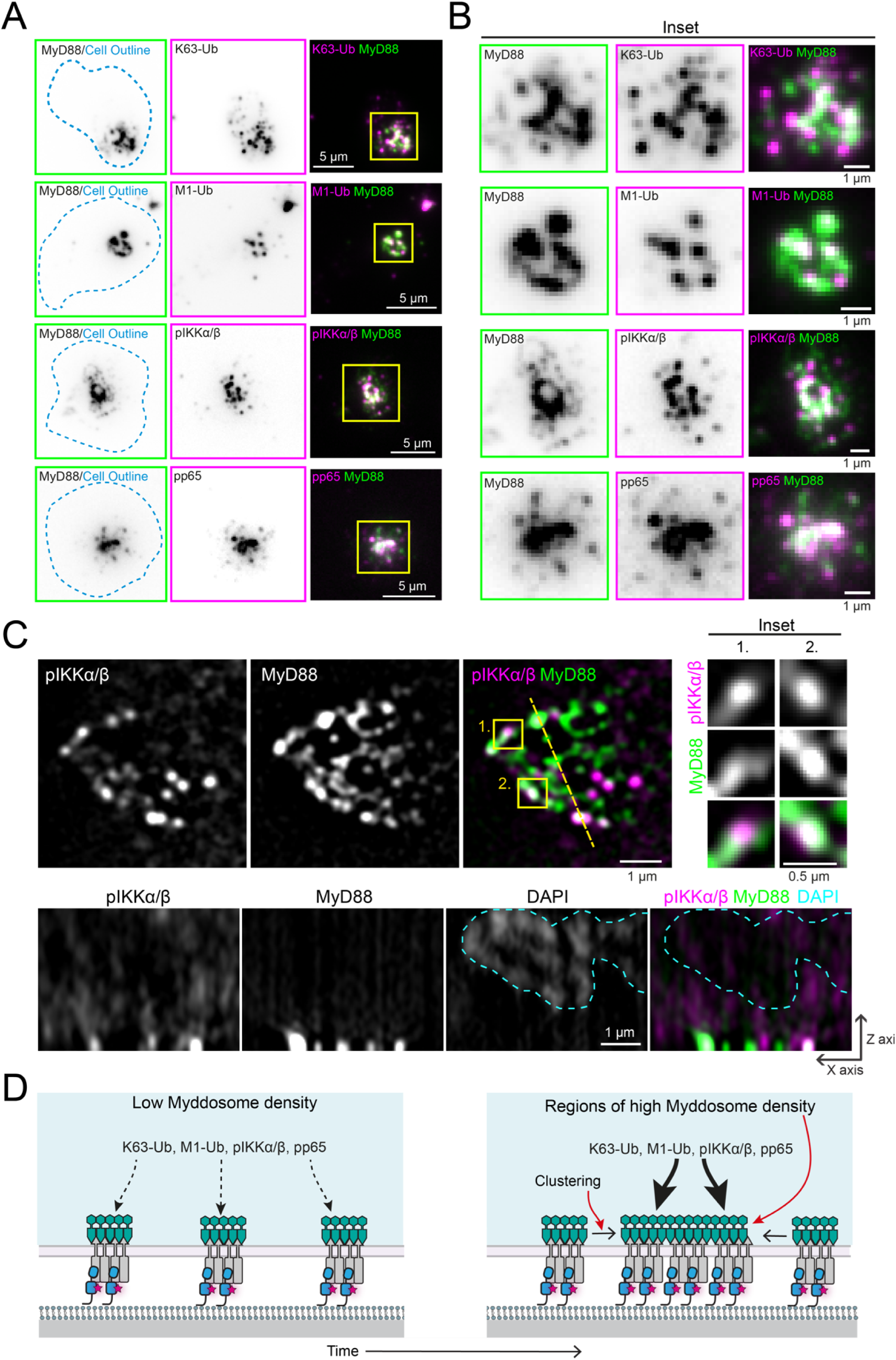
**Myddosomes colocalize with a NF-kB signalosome composed of K63-Ub and M1-Ub chains and phospho-IKK and phospho-p65** (A) TIRF images of fixed EL4-MyD88-GFP cells and stained with antibodies against K63-Ub, M1-Ub, pIKK and pp65. Scale bar, 5 µm. (B) A magnified view of the large patch-like Myddosome clusters from the highlighted region of interest in panel A (yellow box on merge images). Scale bar, 1 µm. (C) Structured illumination microscopy images of Myddosome clusters stained with anti-pIKK. Top row right, insets show detail of Myddosome staining with anti-pIKK. Inset taken from regions of interest overlaid the merge image (yellow boxes 1 and 2). Bottom row, x-z view slice taken from yellow line overlaid on the merge image (top row). Myddosome and pIKK staining localize the cell-SLB interface. Scale bar, 1 µm. (D) Schematic showing working model for how Myddosome clustering could enhance the generation of K63/M1-Ub, pIKK and pp65 and a NF-kB signalosome. We hypothesize that the Myddosome clustering creates regions with a high density of complexes, and this will lead to enhanced production of signaling intermediates such as K63-Ub and M1-Ub chains, pIKK and pp65.

We used structured illumination microscopy (SIM) to image the spatial organization of pIKK and MyD88 puncta with higher resolution and within the entire cellular volume (Fig. 2C). Consistent with our TIRF studies, we found that pIKK punctate structures colocalized with MyD88-GFP puncta at the cell surface (Fig. 2C). In some instances, SIM revealed that pIKK puncta partially overlapped or were adjacent to MyD88-GFP puncta (inset, Fig. 2C). Z-stack analysis revealed that pIKK puncta localized to the cell-bilayer interface, and were rarely found deeper within the cytosol (x-z view, Fig. 2C). In summary, Myddosomes function as a scaffold and a focal point for the formation of an NF-kB signalosome that is composed of K63/M1-Ub chains and phosphorylated IKK complex and NF-kB subunits.

### The degree of Myddosome clustering displays a linear relationship with NF-kB signalosome size

Diminished RelA nuclear translocation on nanopatterned grids (Fig. 1H) suggested a connection between the spatial organization of Myddosomes and the degree of NF-kB activation. We wondered if this connection is because multi-Myddosome clusters, as compared to single Myddosomes, function as high efficiency scaffolds and focal points for enhanced NF-kB signalosome assembly (Fig. 2D). To test this hypothesis we used TIRF microscopy to analyze the spatial relationship between Myddosome density and accumulation of pp65 and pIKK. A scatter plot of MyD88 puncta intensity versus pp65 or pIKK staining intensity (Fig. 3A-B) shows a linear relationship (R = 0.64 and 0.71 from pp65 and pIKK, Fig. 3A-B, S3A-B). Thus brighter MyD88 puncta, which most likely corresponded to clusters of multiple Myddosome complexes (Fig. 2D), had increased levels of pIKK and pp65 (see insets, Fig. 3A-B for example). Therefore, the signaling output of Myddosomes increases proportionally with the degree of clustering: the higher the density of complexes within a Myddosome cluster, the greater the intensity of pIKK and pp65.

**Figure 3.**
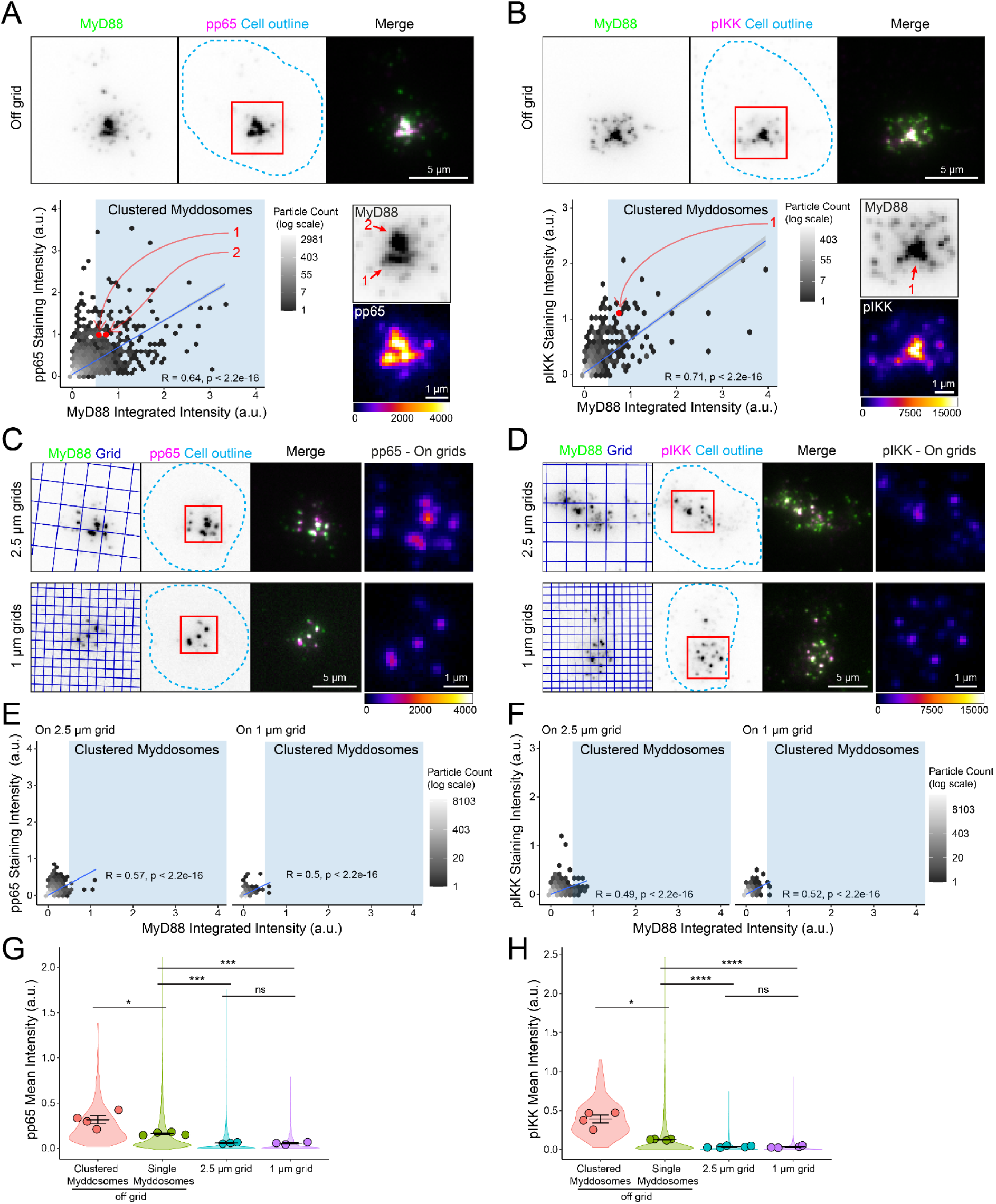
**Comparison of pIKK and pp65 antibody staining at Myddosomes assembled off and on nanopatterned grids.** (**A and B**) Top, TIRF images of fixed EL4-MyD88-GFP cells incubated with IL-1 functionalized SLBs for 30 minutes, and stained with antibodies against pp65 (A) or pIKK (B). Scale bar, 5 μm. Region of interest (red box, merge image) shows an example of MyD88-GFP puncta that colocalizes with pp65 (A) or pIKK (B) puncta. Bottom, 2D histograms of the distribution of MyD88 puncta intensity and associated pp65 (A) or pIKK (B) staining intensity. Linear fit is shown as a blue line superimposed on 2D histograms (Pearson correlation coefficient, R, of linear fit labeled on 2D histograms). Blue shaded regions on scatter plot high MyD88 puncta classified as clustered Myddosomes. Bottom right, zoomed images of the region of interest (red box overlaid merge image, top) show MyD88-GFP channel and associated pp65 (A) and pIKK (B) channel (pp65/pIKK images are displayed with Fire LUT). Red data points on the 2D histogram are from indicated puncta in the MyD88-GFP image (numbered red arrows). Scale bar, 1 µm. (**C and D**) TIRF images of fixed EL4-MyD88-GFP cells incubated with partitioned IL-1 functionalized SLBs (2.5 µm top row and 1 µm bottom row) and stained with anti-pp65 (C) or anti-pIKK (D). Region of interest (red box overlaid merge image) shows examples of MyD88-GFP puncta that colocalize with pp65 (A) or pIKK (B) puncta. Scale bar, 5 µm. Far right, zoomed image of pp65 (C) or pIKK (D) puncta (from region of interest overlaid merge image) displayed with Fire LUT. Scale bar, 1 µm. (**E and F**) 2D histogram of MyD88-GFP puncta intensity and associated pp65 (E) or pIKK (F) staining intensity on 2.5 µm and 1 µm grids. Linear fit is shown as a blue line superimposed on 2D histograms (Pearson correlation coefficient, R, of linear fit labeled on 2D histograms). (**G and H**) Quantification of mean pp65 (G) or pIKK (H) staining intensity for puncta classifieds as single or clustered Myddosomes, and MyD88 puncta formed on 2.5 and 1 µm grids. The normalized mean intensity for clusters, single Myddosomes MyD88 puncta on 2.5 and 1 µm grids are the following: for pp65 0.318 ± 0.044, 0.163 ± 0.009, 0.059 ± 0.005 and 0.057 ± 0.008; for pIKK 0.393 ± 0.051, 0.130 ± 0.004, 0.037 ± 0.006 and 0.035 ± 0.007 (a.u., mean ± SEM, mean value states in the order they appear on plot, left to right). Violin plots show the distribution of all segmented MyD88 puncta. Data points superimposed on the violin plots are the averages from independent experiments. p values are *<0.05, ***<0.001, ****<0.0001. Bars represent mean ± SEM (n = 3 - 4 experimental replicates for pp65 with ≥259 puncta measured per replicate for each experimental condition; n = 4 - 5 experimental replicates for pIKK with ≥435 puncta measured per replicate).

We used SLBs formed on 1 and 2.5 µm grids to inhibit the formation of Myddosome clusters (Fig. 1F) and assayed how this impacted pp65 and pIKK staining. We found that cells on grids still assembled MyD88 puncta that colocalized with pp65 and pIKK (Fig. 3C-D, S3A-B). Scatter plot analysis of MyD88 puncta assembled revealed that, similar to off grid (Fig. 3A-B), there was a linear relationship between puncta intensity and associated pp65/pIKK staining (Fig. 3E-F). Similar to above (Fig. 3A-B), this data suggests a linear relationship between the density of Myddosome complexes and pIKK and pp65 production. However, restricting the degree of clustering proportionally reduced pp65 and pIKK production.

We compared the mean pp65/pIKK intensity of puncta classified as single or clustered Myddosomes with puncta formed on 2.5 and 1 µm grids (methods and Fig. S3C-D). We found that Myddosome clusters had a 5-fold and 10-fold greater mean pp65 and pIKK staining intensity compared to MyD88 puncta on 1 and 2.5 µm grids that were most likely single complexes (Fig. 3G-H). Interestingly, single Myddosomes off grid had statistically greater mean pp65/pIKK intensity than single Myddosomes formed on grids. We noticed that off grid some of these single Myddosomes with high pp65/pIKK staining intensity were closely associated with Myddosome clusters, this suggested Myddosome clusters could enhance signaling output for adjacent complexes. The greater concentrations of pp65 and pIKK at Myddosome clusters, suggests they are hot spots for NF-kB signaling, and that the localized production of these outputs is proportional to the degree of complex density within clusters.

We examined the relationship between MyD88-GFP puncta intensity and staining with antibodies against K63-Ub and M1-Ub chains (Fig. 4A-D, S4A-B). Like above, we found a linear correlation between MyD88 puncta intensity and K63/M1-Ub staining intensity (R = 0.75 and 0.73 for K63-Ub and M1-Ub staining intensity, scatter plot, Fig. 4A-B). MyD88 puncta that formed on 1 and 2.5 µm grids still colocalized with punctate K63/M1-Ub structures, but overall had lower staining intensities (Fig. 4C-D). However, similar to off grid, there was a correlation between MyD88 puncta and K63/M1-Ub intensity on grids (Fig, 4C-F). We found that Myddosome clusters had a 4 fold greater mean K63-Ub intensity and a 3 fold greater mean M1-Ub intensity than single Myddosomes and puncta on 1 and 2.5 µm grids (Fig. 4G-H, S4C-D). Similar to pp65 and pIKK staining, single Myddosomes off grid had statistically greater K63-Ub staining intensity than on grid Myddosomes (Fig. 4G). In contrast, single Myddosomes on and off grids had no statistical difference in mean M1-Ub staining intensity (Fig. 4H). Therefore, similar to pp65 and pIKK, the localized generation of K63-Ub and M1-Ub at the cell surface increases proportionally with the nano-scale density of Myddosomes. However, the generation of M1-Ub was specifically enhanced at Myddosome clusters.

**Figure 4.**
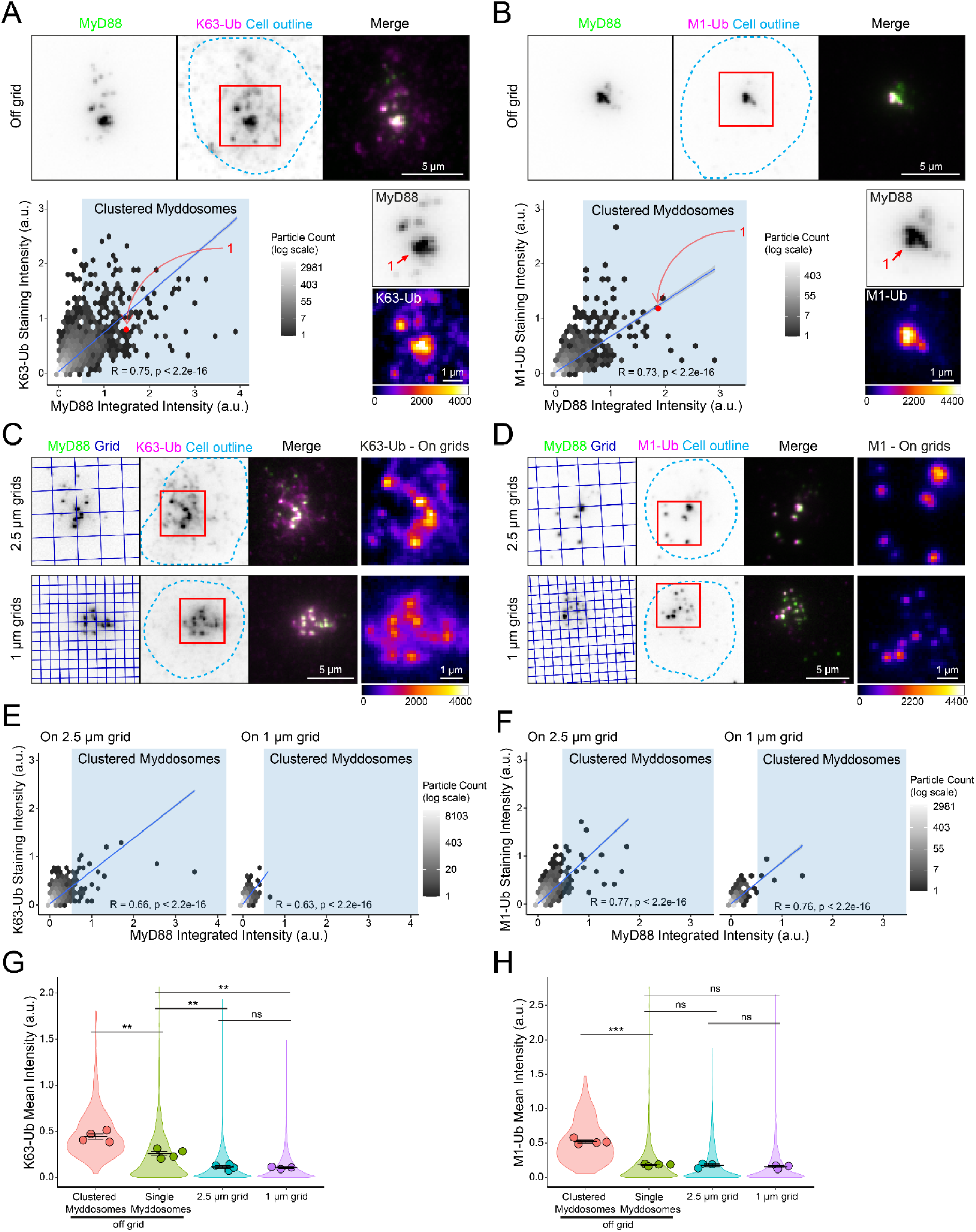
**Comparison of Myddosome K63-Ub and M1-Ub antibody staining on and off nanopatterned grids.** (**A and B**) Top, TIRF images of fixed EL4-MyD88-GFP cells incubated with IL-1 functionalized SLBs for 30 minutes, and stained with anti-K63-Ub (A) or anti-M1-Ub (B). Scale bar, 5 µm. Region of interest (red box, merge image) shows an example of MyD88-GFP puncta that is colocalized with K63-Ub (A) or M1-Ub (B) puncta. Bottom, 2D histograms of the distribution of MyD88 puncta intensity and associated K63-Ub (A) or M1-Ub (B) staining intensity. Linear fit is shown as a blue line superimposed on 2D histograms (Pearson correlation coefficient, R, of linear fit labeled on 2D histograms). Blue shaded region on scatter plot high MyD88 puncta classified as clustered Myddosomes. Bottom right, zoomed images of the region of interest (red box overlaid merge image, top) show MyD88-GFP channel and associated K63-Ub (A) and M1-Ub (B) channel (K63/M1-Ub images are displayed with Fire LUT). Red data points on the 2D histogram are from indicated puncta in the MyD88-GFP image (numbered red arrows). Scale bar, 1 µm. (**C and D**) TIRF images of fixed EL4-MyD88-GFP cells incubated with partitioned IL-1 functionalized SLBs (2.5 µm top row and 1 µm bottom row grids) and stained with anti-K63-Ub (C) or anti-M1-Ub (D). Region of interest (red box overlaid merge image) shows an example of MyD88-GFP puncta that colocalize with K63-Ub (A) or M1-Ub (B) puncta. Scale bar, 5 µm. Far right, zoomed image of K63-Ub (C) or M1-Ub (D) puncta (from region of interest overlaid merge image) displayed with Fire LUT. Scale bar, 1 µm. (**E and F**) 2D histograms of the distribution of MyD88 puncta intensity and associated K63-Ub (E) or M1-Ub (F) staining intensity on 2.5 µm and 1 µm grids. Linear fit is shown as a blue line superimposed on 2D histograms (Pearson correlation coefficient of linear fit labeled on 2D histograms). (**G and H**) Quantification of mean K63-Ub (G) or M1-Ub (H) staining intensity for puncta classifieds as single or clustered Myddosomes, and MyD88 puncta formed on 2.5 and 1 µm grids. The normalized mean intensity for clustered, single Myddosomes, and MyD88 puncta on 2.5 and 1 µm grids are the following: for K63-Ub 0.444 ± 0.030, 0.257 ± 0.025, 0.113 ± 0.015 and 0.104 ± 0.008; for M1-Ub 0.520 ± 0.020, 0.183 ± 0.008, 0.174 ± 0.022 and 0.153 ± 0.016 (a.u., mean ± SEM, mean value states in the order they appear on plot, left to right). Violin plots show the distribution of all segmented MyD88 puncta. Data points superimposed on the violin plots are the averages from independent experiments. p values are **<0.01, ***<0.001. Bars represent mean ± SEM (n = 3 - 4 experimental replicates for K63 with ≥484 puncta measured per replicate for each experimental condition; n = 3 - 4 experimental replicates for M1 with ≥338 puncta measured per replicate).

In summary, we found that signaling outputs such as pp65, pIKK and K63/M1-Ub colocalize with single and clustered Myddosome complexes. This suggests that a single Myddosome can activate NF-kB signalosome formation and this is an invariant signaling output encoded by Myddosome assembly. However, we found Myddosome clusters colocalized with >3 fold larger NF-kB signalosomes defined by greater amounts of pp65, pIKK and K63/M1-Ub. Furthermore, the degree of clustering led to a proportional increase in signalosome size and the robust incorporation of M1-Ub into this NF-kB signalosome. This suggests that the spatial organization of the Myddosome encodes an analog signaling response.

### Larger Myddosome clusters have enhanced TRAF6 and LUBAC recruitment

Having shown that Myddosomes can generate invariant and proportional outputs, we examined how these outputs arose from the dynamics of Myddosome formation, clustering and NF-kB signalosome assembly. Our data (Fig. 2-4), along with published studies (Tarantino et al. 2014; Du et al. 2022), suggest that NF-kB activation occurs in condensate cellular compartments that contain K63-Ub/M1-Ub chains. We generated two CRISPR double knock-in EL4 cell lines that expressed MyD88-GFP and either the K63-Ub E3 ligase TRAF6 or the M1-Ub E3 ligase LUBAC subunit HOIL1 labeled with the mScarlet (Fig. S5). When we imaged these cell lines we found that a subset MyD88-GFP puncta recruited mScarlet-TRAF6 (Fig. 5A) or mScalet-HOIL1 (Fig. 5B). We found that TRAF6 or HOIL1 appeared after the formation of MyD88 puncta (Fig. 5A-B, Movie 3-4). Both MyD88-GFP and mScarlet-TRAF6 or mScarlet-HOIL1 puncta were initially dim and grew in intensity (Fig. 5A-B). In some instances, we observed that TRAF6 was transiently recruited to Myddosomes, with this transient TRAF6 recruitment often preceding the stable association of TRAF6 with MyD88 (Fig. S6A). This dynamic suggests that Myddosome served as a scaffold for the nucleation of TRAF6 assemblies (Yin et al. 2009). In summary, we found that Myddosomes recruit and assemble punctate structures of the ubiquitin ligases TRAF6 and LUBAC. The molecular dynamics of MyD88 and the E3 ligases TRAF6 or LUBAC are consistent with Myddosomes functioning as an inducible scaffold and focal point for the activation of K63/M1-Ub generation.

**Figure 5.**
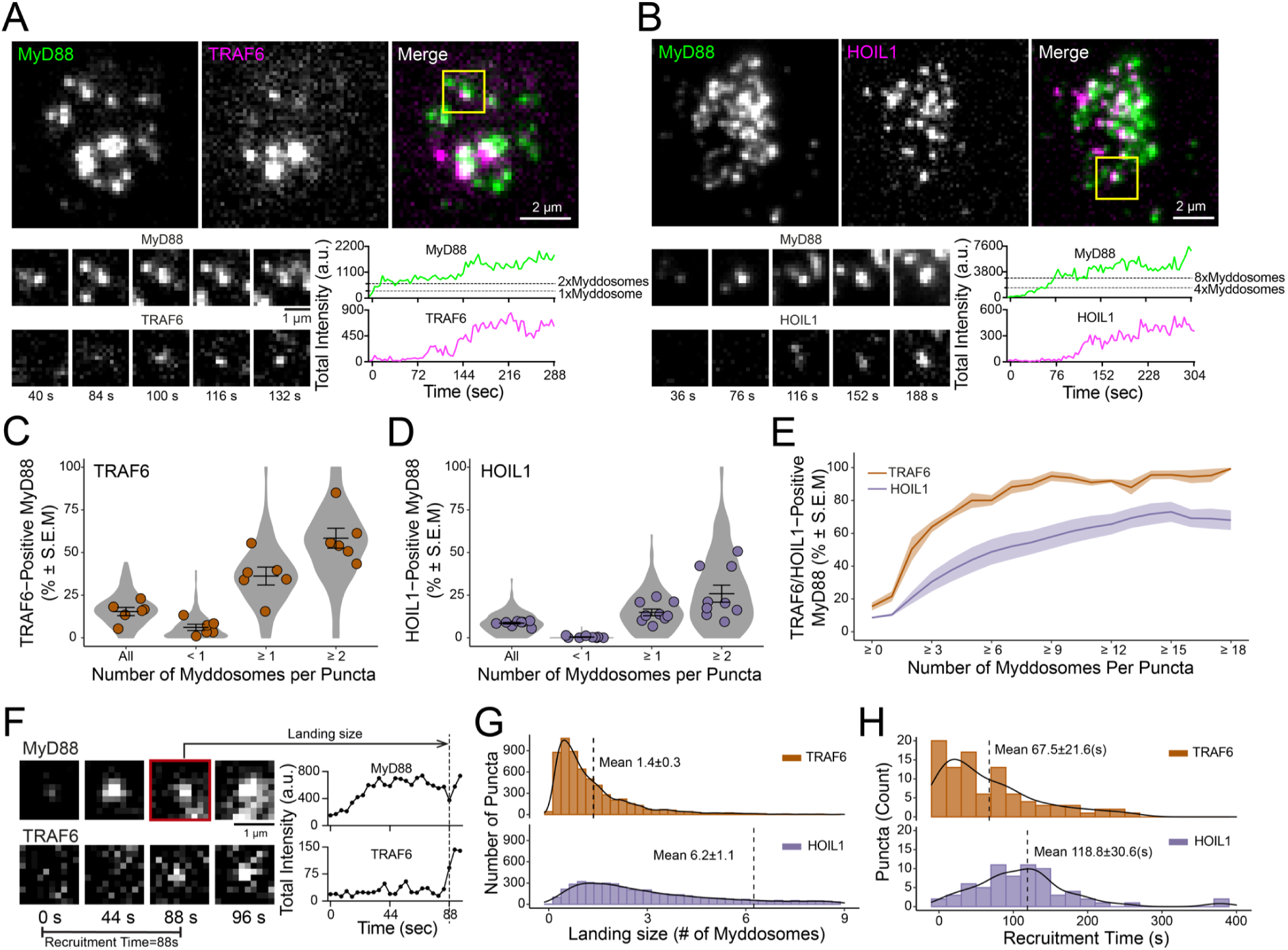
**E3 ligases TRAF6 and HOIL1 are recruited to Myddosomes.** (**A and B**) Top: TIRF images of MyD88-GFP and mScarlet-TRAF6 (A) or mScarlet-HOIL1 (B). Region of interest (yellow box, merge image) shows an example of a MyD88-GFP puncta colocalized with mScarlet-TRAF6 (A) or mScalet-HOIL1 (B). Bottom: Time-series TIRF images from the region of interest (left) and fluorescence intensity times series (right) of MyD88 and TRAF6 (A) or HOIL1 (B). (**C and D**) Quantifications of percentage of MyD88-GFP puncta that colocalized with TRAF6 (C) or HOIL1 (D) grouped for all puncta, puncta containing <1x, ≥1x and ≥2x Myddosome complexes. The percentages for TRAF6 (C) in these groups are 15.4 ± 2.4%, 6.1 ± 1.8%, 36.2 ± 5.2% and 58.4 ± 5.9%, respectively (mean ± SEM). The percentages for HOIL1 (D) in these groups are 8.6 ± 0.5%, 0.4 ± 0.2%, 14.9 ± 1.9% and 25.9 ± 5.0%, respectively (mean ± SEM). Violin plots indicate the distribution of individual cell measurements. Colored dots superimposed on the violin plots are the averages from independent experiments. Bars represent mean ± SEM (n = 6 experimental replicates for TRAF6, with 17-48 cells measured per replicate; n = 9 experimental replicates for HOIL1, with 9-46 cells measured per replicate). (**E**)Quantifications of the percentage of TRAF6-or HOIL1-positive MyD88 puncta as a function of Myddosome numbers per MyD88 puncta. With greater Myddosome numbers per puncta, the percentage of MyD88 colocalized with TRAF6 or HOIL1 increases. The line represents the average across replicates and the shaded area represents SEM. (**F**) Analysis of TRAF6 and HOIL1 landing size and recruitment time to MyD88 puncta. Left, time series TIRF images showing MyD88-GFP puncta nucleation and the appearance of TRAF6. Right, the associated fluorescence intensity time trace for the time series shown. Recruitment time is defined as the time interval from Myddosome nucleation (e.g., time = 0 s when MyD88-GFP puncta appears) to the appearance of a TRAF6 or HOIL1 puncta. Landing size is defined as the fluorescent intensity of the MyD88 puncta at the time when TRAF6 or HOIL1 appears (indicated on the fluorescent intensity trace with arrow). (**G**) Histogram of landing size of MyD88 puncta, expressed as number of Myddosome complexes per puncta, for TRAF6 (top, n = 6015 recruitment events from 183 cells) and HOIL1 (bottom, n = 5562 recruitment events from 212 cells) recruitment. Histogram is overlaid with a density plot of the distribution. Black horizontal lines on the histograms denote the average landing size (mean ± SD). (**H**) Histogram for the recruitment time of TRAF6 (top, n = 94 recruitment events from 4 cells) and HOIL1 (bottom, n = 69 recruitment events from 4 cells) overlaid with the density plot of the distribution. Black horizontal lines on the histograms denote the average recruitment time (mean ± SD).

We asked whether MyD88-GFP puncta clustering and lifetime enhanced TRAF6 recruitment. We observed that TRAF6 positive MyD88 puncta had an average size of 10.4x MyD88s (Fig. S6B-C). Based on structural studies of 6x MyD88s per Myddosome (S.-C. Lin, Lo, and Wu 2010), the average MyD88 copy number in TRAF6 positive puncta suggested they contain on average one or more Myddosome complexes. In total 15.4 ± 2.4% of MyD88 puncta colocalize with TRAF6 (mean ± SEM, from 6 replicates, Fig. 5C, All); however, when we normalized MyD88 puncta intensity to the number Myddosomes per puncta (see Methods, and Fig. S6B), we found that 58.4 ± 5.9% of Myddosome clusters were TRAF6 positive (Fig. 5C, ≥2). In comparison, the percentage of TRAF6 positive puncta was 6.1 ± 1.8% and 36.2 ± 5.2% for puncta containing <1 or ≥1 Myddsome complexes (mean ± SEM, from 6 replicates, Fig. 5C and S6E). We analyzed the relationship between lifetime and TRAF6 recruitment. Using a threshold of 50 s to define long lived MyD88 puncta, we found that 34.6 ± 3.5% MyD88 with lifetimes ≥50s colocalized with TRAF6 versus 7.1 ± 1.4% of puncta with lifetime <50s (mean ± SEM, from 6 replicates, Fig. S6D). In summary, stable Myddosome clusters are more likely to recruit TRAF6.

We applied the same analysis to investigate HOIL1 recruitment. We found that HOIL1 positive MyD88 puncta were greater in size than non-colocalized puncta (47.4 MyD88s for positive puncta vs 11.6 MyD88s for negative puncta, Fig. S6F). 19.3 ± 1.3% of MyD88 puncta with a lifetime ≥ 50s colocalized with HOIL1. In comparison only 2.5 ± 0.2% of MyD88 puncta with lifetime <50s colocalized with HOIL1 (Fig. S6G). In summary, like observed for TRAF6, MyD88 puncta that recruit HOIL1 were more likely to be larger puncta with longer lifetimes. We investigated the role of Myddosome clustering in HOIL1 recruitment. We found that on average 8.6 ± 0.5% of all MyD88-GFP puncta colocalized with HOIL1 (mean ± SEM%, from 9 replicates, Fig. 5D, S6H). When we quantified multi-complex puncta containing ≥1 or ≥2 Myddosomes, we found that the percent of HOIL1 positive recruitment increased to 14.9 ± 1.9% and 25.9 ± 5.0% respectively (mean ± SEM, Fig. 5D).

Finally, we plotted the cumulative distribution of the percent of MyD88 puncta that colocalized with TRAF6 or HOIL1 as a function of Myddosomes per MyD88 puncta (Fig. 5E). We found that the percent of TRAF6 and HOIL1 colocalized puncta increased as the number of Myddosomes per MyD88-GFP puncta increased (Fig. 5E, S6I, S6J). Therefore, Myddosomes organized into clusters have a greater probability of recruiting E3 ubiquitin ligases TRAF6 and HOIL1, suggesting a mechanistic basis for why K63-Ub/M1-Ub scales with the density of Myddosome complexes within clusters (Fig. 4).

### Myddosome clustering triggers the sequential recruitment of TRAF6 and LUBAC

If clustering is a driver for TRAF6 and HOIL1 recruitment, we expect that formation of clusters would precede the recruitment of both ligases. Therefore, we asked whether the formation of Myddosome clusters occurs before or after the recruitment of TRAF6 and HOIL1. We analyzed the size of Myddosomes at the time point when TRAF6 and HOIL1 are recruited. We defined this time point as the TRAF6/HOIL1 landing size (Fig. 5F) and quantified the number of Myddosomes per puncta at this time point. We found that the average landing size for TRAF6 was 1.4 ± 0.3 Myddosome complexes and for HOIL1 the landing size was 6.2 ± 1.1 Myddosome complexes per puncta (Fig. 5G). This suggests that on average, the formation of Myddosome clusters precedes the recruitment of TRAF6 and LUBAC.

We analyzed the recruitment time of TRAF6 and HOIL1, which we defined as the time interval from the nucleation of a MyD88 puncta to the recruitment of mScarlet-TRAF6 or mScarlet-HOIL1 (Fig. 5H). We found that the average recruitment time for TRAF6 was 67.5 ± 21.6 secs (mean ± SD, Fig. 5H). In contrast HOIL1 had an average recruitment time of 118.8 ± 30.6 secs (mean ± SD, Fig. 5H). We conclude that TRAF6 and HOIL1 are recruited to clusters of Myddosomes and that the recruitment of these two ubiquitin ligases is staggered temporally: TRAF6 is recruited first followed by HOIL1. In contrast to TRAF6, HOIL1 is recruited to puncta composed of a greater density of Myddosome complexes. In conclusion, Myddosome signaling outputs are kinetically controlled by spatial organization. Specifically the analog production of signaling outputs are encoded by Myddosome density within clusters as this regulates the probability of TRAF6 and HOIL1 recruitment.

### Perturbing Myddosome clustering diminishes TRAF6 recruitment

We set out to assay how the combination of nanopattern grids and ligand density affected Myddosome clustering and TRAF6/HOIL1 recruitment. If clustering regulated the probability of TRAF6/LUBAC recruitment (Fig. 5E, G) and this was the basis of digital and analog Myddosome signaling outputs, we reasoned inhibiting clustering and isolating single complexes should reduce the recruitment of both E3 ligases. We predicted that increasing IL-1 density within individual 1 µm^2^ corrals would restore TRAF6/HOIL1 recruitment, as single corrals would contain a sufficient number of IL-1 to trigger the assembly of multiple Myddosomes that could coalesce into clusters.

We characterized the formation of Myddosome clusters in EL4-MyD88-GFP/mScarlet-TRAF6 cells stimulated by SLBs functionalized with 1 IL-1/ µm^2^ on and off 1 µm grids. We found that only 1.2 ± 0.7% of MyD88 puncta were classified as clusters in EL4-MyD88-GFP/mScarlet-TRAF6 cells stimulated with 1 µm^2^ gridded SLBs at this ligand densities (mean ± SEM, Fig. S7A). In contrast, we measured that 2.7 ± 1.4% of MyD88 puncta were clusters on SLB formed off nanopatterned grids (mean ± SEM, Fig. S7A). As observed previously (Fig. 1F), we found that MyD88-GFP puncta size in cells stimulated with 1 µm grids are smaller than those stimulated with SLBs off grid (a mean of 2.2 versus 1.7 MyD88s per puncta off and on grids, Fig. S7A). Live cell imaging and kymograph analysis at this ligand density showed that MyD88/TRAF6 puncta coalescence and clustered (Fig. 6A, Movie 5). Similar to above (Fig. 5A), these dynamic MyD88 puncta recruited mScarlet-TRAF6. In contrast, we observed that MyD88-GFP puncta on 1 µm grids did not coalesce and cluster (kymograph, Fig. 6B). However a portion of these puncta still recruited mScarlet-TRAF6 (Fig. 6B, Movie 5). Analysis revealed a two fold difference off and on grids in the frequency of TRAF6 recruitment at 1 IL-1/µm^2^ (16.2 ± 2.5% vs. 6.1 ± 1.5% TRAF6 positive MyD88 puncta per cells off and on grids respectively, mean ± SEM, Fig. 6C and Fig. S7C).

**Figure 6.**
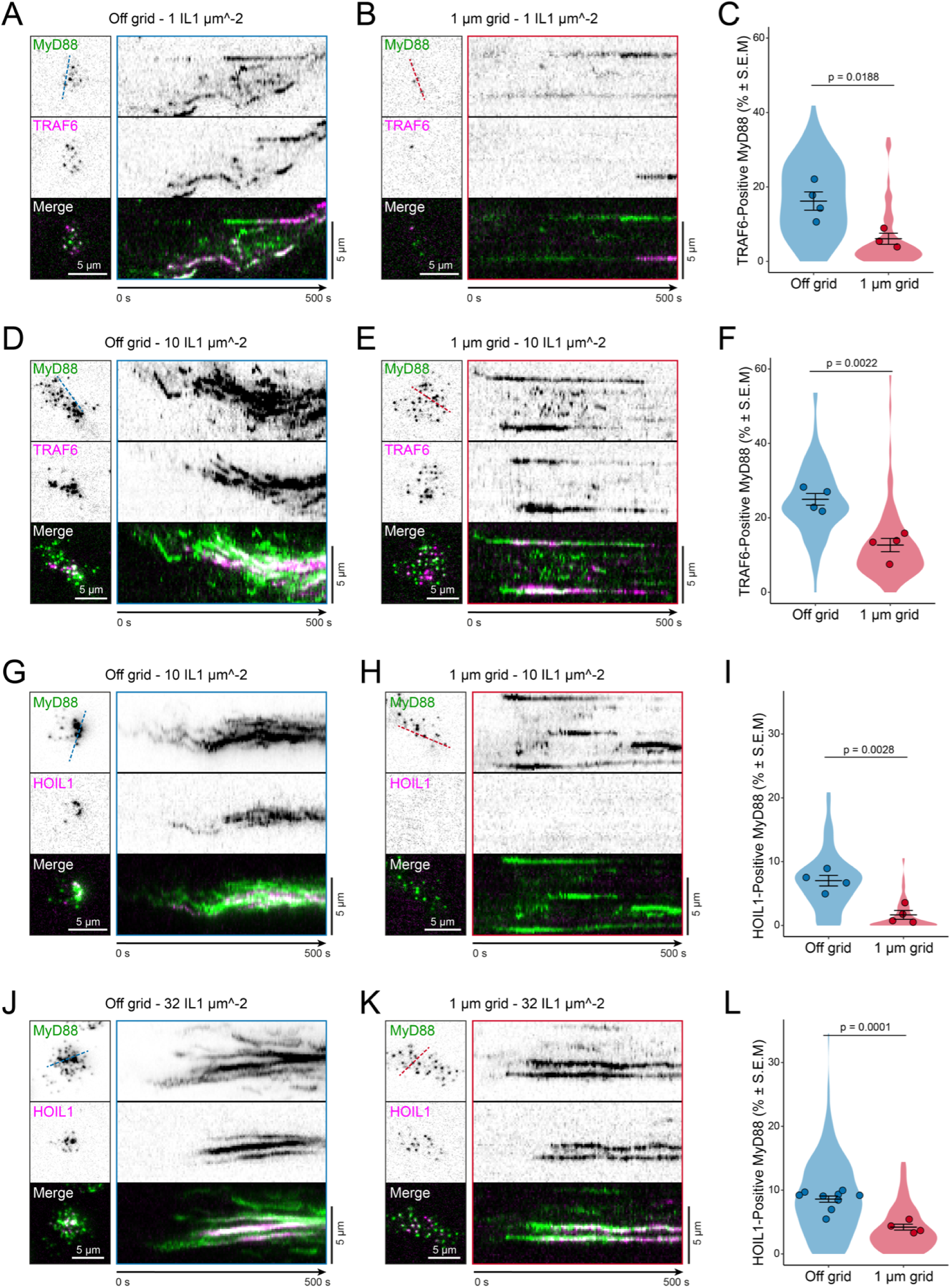
**Extracellular barriers that restrict Myddosome clustering reduce TRAF6 and HOIL1 recruitment.** (**A and B**) TIRF images of EL4 cells expressing MyD88-GFP and mScarlet-TRAF6 stimulated on IL-1 functionalized SLBs at a ligand density of 1 IL-1/µm^2^ off grids (A) or on 1 µm grids (B). Kymographs derived from dashed lines overlaid TIRF images (left panel). Scale bar, 5 µm. (C) Quantification of percentage of MyD88-GFP puncta that colocalized with TRAF6 off grids and on 1 µm grids at a ligand density of 1 IL-1/µm^2^, and the percentages are 16.2 ± 2.5% and 6.1 ± 1.5%, respectively (mean ± SEM). Violin plots indicate the distribution of individual cell measurements. Colored dots superimposed on the violin plots are the averages from independent experiments. Bars represent mean ± SEM (n = 4 experimental replicates off grids, with 18-27 cells measured per replicate; n = 3 experimental replicates on 1 µm grids, with 17-34 cells measured per replicate). (**D and E**) TIRF images of EL4 cells expressing MyD88-GFP and mScarlet-TRAF6 stimulated on IL-1 functionalized SLBs at a ligand density of 10 IL-1 per µm^2^ off grids (D) or on 1 µm grids (E). Kymographs derived from dashed lines overlaid TIRF images (left panel). Scale bar, 5 µm. (**F**) Quantification of percentage of MyD88-GFP puncta that colocalized with TRAF6 off grids and on 1 µm grids at a ligand density of 10 IL-1/µm^2^, and the percentages are 25.0 ± 1.6% and 12.7 ± 1.8%, respectively (mean ± SEM). Violin plots indicate the distribution of individual cell measurements. Colored dots superimposed on the violin plots are the averages from independent experiments. Bars represent mean ± SEM (n = 4 experimental replicates off grids, with 9-33 cells measured per replicate; n = 4 experimental replicates on 1 µm grids, with 17-33 cells measured per replicate). (**G and H**) TIRF images of EL4 cells expressing MyD88-GFP and mScarlet-HOIL1 stimulated on IL-1 functionalized SLBs at a ligand density of 10 IL-1/µm^2^ off grids (G) or on 1 µm grids (H). Kymographs derived from dashed lines overlaid TIRF images (left panel). Scale bar, 5 µm. (**I**) Quantification of percentage of MyD88-GFP puncta that colocalized with HOIL1 off grids and on 1 µm grids at a ligand density of 10 IL-1/µm^2^, and the percentages are 7.0 ± 0.8% and 1.7 ± 0.7%, respectively (mean ± SEM). Violin plots indicate the distribution of individual cell measurements. Colored dots superimposed on the violin plots are the averages from independent experiments. Bars represent mean ± SEM (n = 4 experimental replicates off grids, with 16-37 cells measured per replicate; n = 4 experimental replicates on 1 µm grids, with 25-50 cells measured per replicate). (**J and K**) TIRF images of EL4 cells expressing MyD88-GFP and mScarlet-HOIL1 stimulated on IL-1 functionalized SLBs at a ligand density of 32 IL-1/µm^2^ off grids (J) or on 1 µm grids (K). Kymographs derived from dashed lines overlaid TIRF images (left panel). Scale bar, 5 µm. (**L**) Quantification of percentage of MyD88-GFP puncta that colocalized with HOIL1 off grids and on 1 µm grids at a ligand density of 32 IL-1/µm^2^, and the percentages are 8.6 ± 0.5% and 4.2 ± 0.5%, respectively (mean ± SEM). Violin plots indicate the distribution of individual cell measurements. Colored dots superimposed on the violin plots are the averages from independent experiments. Bars represent mean ± SEM (n = 9 experimental replicates off grids, with 9-46 cells measured per replicate; n = 4 experimental replicates on 1 µm grids, with 15-45 cells measured per replicate).

We assayed TRAF6 recruitment at a higher ligand density of 10 IL-1/µm^2^, to test if this increases the percentage of TRAF6 positive MyD88 puncta. At 10 IL-1/µm^2^ the size of MyD88 puncta on 1 µm grids was still smaller than puncta off grid but larger than at 1 IL-1/µm^2^ (average size of 1.7 versus 2.6 MyD88s on grids at 1 and 10 IL-1/µm^2^ respectively, Fig. S7A-B), and the percentage of Myddosome clusters increased by four fold (4.7% versus 1.2% of puncta classified as clusters at 10 and 1 IL-1/µm^2^, mean ± SEM, Fig. S7A-B). Importantly, compared to 1 IL-1/µm^2^, the percentage of TRAF6 positive Myddosome increased on 1 µm grids functionalized with 10 IL-1/µm^2^ (12.7 ± 1.8% versus 6.1 ± 1.5% TRAF6 positive MyD88 puncta per cells at 10 and 1 IL-1/µm^2^, Fig. 6C-F and Fig. S7C, Movie 6). Thus, increasing the number of IL-1 per 1 µm^2^ corral can rescue the perturbation of TRAF6 recruitment. In summary, this data reveals that single Myddosomes can trigger a TRAF6 response, consistent with an invariant signaling output. However, Myddosome clusters have increased probability of TRAF6 recruitment, suggesting a mechanism for how clusters can generate a proportionally greater signaling output than single complexes.

### Myddosome clustering triggers enhanced LUBAC recruitment

We examined HOIL1 recruitment off and on 1 µm grids at two SLB ligand densities. We found that 43% and 50% of MyD88 puncta could be classified as clusters composed of multiple Myddosome complexes in MyD88-GFP/mScarlet-HOIL1 cells stimulated by SLB off grids at 10 or 32 IL-1/µm^2^ respectively (Fig. S8A-B). In contrast, on 1 µm grids at 10 IL-1/µm^2^ 3% of MyD88 puncta were clusters (Fig. S8A); however, 13% of puncta were Myddosome clusters at 32 IL-1/µm^2^ (Fig. S8B). Thus by manipulating ligand density on 1 µm grids we could inhibit and restore the formation of Myddosome clusters. At a ligand density of 10 IL-1/µm^2^, we observed the dynamic coalescence and clustering of MyD88 puncta (Fig. 6G, Movie 7). As observed previously (Fig. 5B), mScarlet-HOIL1 was recruited to these large clusters of Myddosomes. However, on 1 µm grids at 10 IL-1/µm^2^, we observed that <2% puncta recruited HOIL1 (7.0 ± 0.8% versus 1.7 ± 0.7% puncta recruited HOIL1 off grid versus on grid, see Fig. 6H, 6I and Fig. S8C-D, Movie 7). We analyzed the density of Myddosomes per HOIL1 positive MyD88 puncta on and off grids, and found that the average number of Myddosomes per puncta at the point of HOIL1 recruitment was 4.4 off grid and 0.7 on 1 µm grids (Fig. S8E). This data suggests that single complexes can recruit HOIL1, but the probability was >4 fold lower than at Myddosome clusters.

We assayed HOIL1 recruitment at 32 IL-1/µm^2^. Off grid, we observed robust HOIL1 recruitment, as expected (Fig. 6J, Movie 8). On 1 µm g rids at this higher ligand density, we found that MyD88 puncta readily recruited HOIL1 (8.6 ± 0.5% versus 4.2 ± 0.5% HOIL1 positive MyD88 puncta off and on 1 µm grids, respectively Fig. 6K,L, also see Fig. S8D, Movie 8). At 32 IL-1/µm^2^ the HOIL1 landing size was 6.2 vs 1.4 Myddosome complexes per puncta off vs on grid respectively (Fig. S8F), suggesting that in both experimental regimes HOIL1 is recruited to Myddosome clusters. We examined the lifetime of TRAF6 and HOIL1 recruitment to single and clustered Myddosomes from on and off grid data (Fig 7A,B). We find that a greater density and number of Myddosome complexes within clusters correlates with a greater lifetime of TRAF6 and HOIL1. We conclude that Myddosome clusters increase the stability of TRAF6 and HOIL1 at Myddosomes, that this is the possible basis for why clusters have increased signaling output. In summary, we can control the association of LUBAC with Myddosome complexes by controlling the formation of clusters and regions of the plasma membrane containing a high density of Myddosomes (Fig. 2D). We conclude that clustering triggers LUBAC recruitment to Myddosomes, and activates M1-Ub production (Fig. 4H). Our results show that Myddosome clustering and the sequential recruitment of TRAF6 and then HOIL1 (Fig. 5H) is a potential mechanism to generate signaling outputs that are proportional to the level of stimulation.

**Figure 7.**
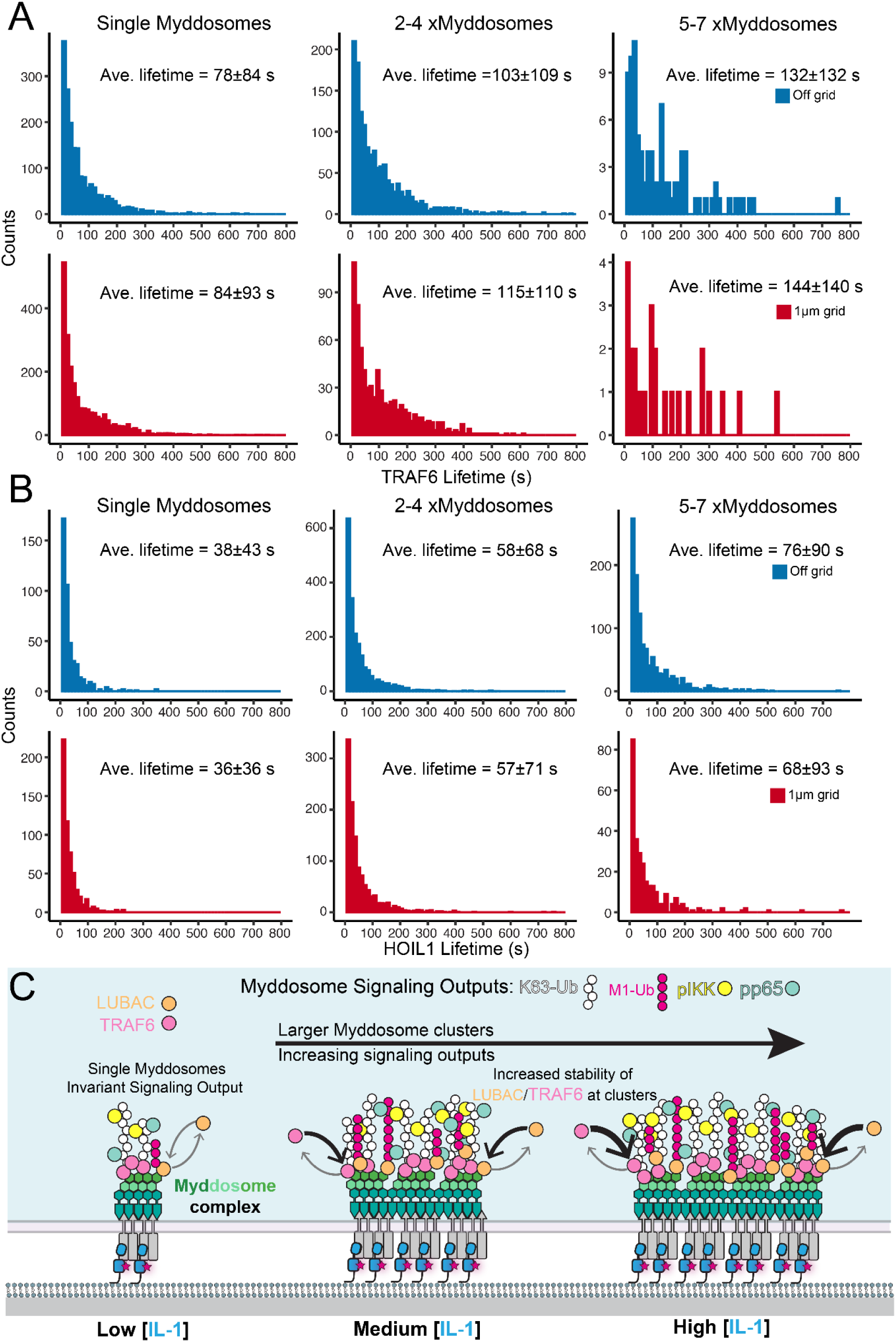
Clustering increases TRAF6 and HOIL1 lifetime at Myddosomes. **(A-B)** Histogram showing the lifetime of TRAF6 and HOIL1 recruitment to single Myddsomes, and clusters containing 2-4 and 5-7 Myddosome complexes, off (blue) and on (red) 1 µm grids. **(C)** Model showing how a single Myddosome has a digital signaling output. However, the amplitude of this output increases proportionally as the density of Myddosome complexes increases within clusters. The increased amplitude of the M1/K63-Ub, pIKK, and p65 signaling output is likely due to the increase in stability of LUBAC and TRAF6 at larger clusters. As Myddosomes are biochemically coupled to extracellular IL-1, this mechanism examines how IL-1 signaling can generate both digital and analog signaling responses that are proportional to the stimulating dose of IL-1.

## Discussion

Here we used high-resolution microscopy to visualize and quantify the signaling output of Myddosomes. Using extracellular nanoscale barriers, we find that Myddosomes are tethered to the cell surface via a direct interaction with the IL-1R bound to extracellular IL-1 (Fig. 1B-C). Myddosomes aggregate into larger cell surface assemblies. The result of this dynamic is the formation of regions on the plasma membrane that contains a high local density of Myddosomes. These regions of high Myddosome density preferentially nucleate the formation of a signalosome composed of the Ubiquitin ligases TRAF6 and LUBAC, K63/M1-Ub chains, and pIKK (Fig. 2). We found these regions also contain pp65, a subunit of the NF-kB complex, suggesting the function of this signalosome is to activate NF-kB (Fig. 2A-B). Single complexes can recruit TRAF6 and HOIL1 (Fig. 5) and promote the productin of K63/M1-Ub, pIKK and pp65 (Fig. 3-4), suggesting NF-kB signalosome formation is a digital output of Myddosomes. However, the probability of nucleating this NF-kB signalosomes is significantly lower at single complexes versus clusters. These results suggest that the spatial organization of Myddosomes can encode responses that are proportional to the amount of IL-1 stimulation.

Previous studies have found that Myddosomes form large aggregate structures after TLR or IL-1 stimulation (Latty et al. 2018; Deliz-Aguirre et al. 2021; Latz et al. 2002). In macrophages stimulated with TLR4 agonist LPS the formation of large Myddosome clusters correlated with higher doses of LPS stimulation, enhanced NF-kB activation and gene expression (Latty et al. 2018). These results are consistent with our finding that inhibiting the formation of Myddosome clusters reduces RelA nuclear translocation (Fig. 1G). Here we find evidence that clusters of Myddosomes enhance NF-kB signaling by increasing the probability of TRAF6 and, in particular, LUBAC recruitment (Fig. 5C-E). We showed that the number of Myddosome clusters increases with higher IL-1 stimulation (Fig. S7A, B and Fig. S8A, B) and that Myddosomes are biochemically coupled to IL-1:IL-1R complexes (Fig. 1). Therefore, Myddosome clustering ‘reads out’ the amount of IL-1 stimulation and converts it into proportional TRAF6 and HOIL1 recruitment (Fig. 6).

How does clustering enhance TRAF6 and HOIL1 recruitment? The Myddosome has a fixed stoichiometry (S.-C. Lin, Lo, and Wu 2010; Motshwene et al. 2009), and with 4x IRAK1 monomers per complex it has a maximum of 12x TRAF6 binding motifs per complex (Ye et al. 2002). Therefore, clustering might be a dynamic mechanism to increase the avidity of TRAF6 binding sites at a focal point on the plasma membrane. We show that single Myddosomes can still recruit TRAF6 to the cell surface (Fig. 6A), although at a lower probability than clusters of Myddosomes. TRAF6 is predicted to form a 2D lattice (Yin et al. 2009), with the trimeric C-terminus making contact with the Myddosome (Ye et al. 2002). Myddosomes clustering might stabilize higher-order assemblies of TRAF6 that promote its ubiquitin ligase activity (Yin et al. 2009). LUBAC component HOIP recognizes K63-Ub (Emmerich et al. 2013) and thus its recruitment depends on the amount of K63-Ub chains. We find less K63-Ub associated with single Myddosomes than clustered Myddosomes (Fig. 4). We also find HOIL1 and M1-Ub are especially sensitive to Myddosome clustering (Fig. 4, 6). Therefore, Myddosome clustering might lead to larger TRAF6 assemblies, enhanced ubiquitin ligase activity, a greater production of K63-Ub, and enhanced HOIL1 recruitment as well as formation of M1-Ub. Thus, the K63-Ub output of TRAF6 will scale proportionally with the density of Myddosomes within clusters.

In conclusion, Myddosomes function as a plasma membrane-associated scaffold that assembles an NF-kB activating signalosome. We show that the spatial density of the Myddosome regulates the assembly and size of this NF-kB activating compartment. This mechanism might explain how the IL-1 signaling pathway can create invariant and proportional NF-kB responses (DeFelice et al. 2019; Son et al. 2021). Other innate immune signaling pathways, such as inflammasomes and STING, use the clustering of signaling complexes to control the formation of specialized signaling compartments (Magupalli et al. 2020; Yu et al. 2021). It is possible clustering is a unifying mechanism across innate immune signaling to transmit switch-like responses and analog information such as the amount and duration of a stimulus. An important future direction is quantifying the spatial organization of other innate immune signaling complexes and how this connects to digital versus analog signaling responses. The approach we establish here that combines live cell microscopy with technologies that enable spatial control of signaling complexes provides a powerful strategy to study how the dynamics of signaling pathways shape signaling outputs.

## Acknowledgements

We thank Luke Lavis (Janelia Research Campus, Ashburn, VA) for providing Janelia Fluor dyes. We acknowledge Olivia Majer (MPIIB) for access and assistance with 3D structured illumination microscopy and for critical reading of the manuscript. We thank Mark Cronan for reading and commenting on an early version of this manuscript. We thank Paulina Dirvanskyte for helpful suggestions in the writing of this paper. We thank Enfu Hui (UCSD) for critical feedback on the manuscript. We thank all Taylor lab members for critical feedback and commenting on the project.

This work was supported by the Max Planck Society.

The authors declare no competing financial interests.

## Author contributions

M.J. Taylor and E. Ziska, purified proteins for labeling membranes. M.J. Taylor, F. Cao, and E. Ziska designed, prepared the HDR and gRNA plasmids and generated the CRISPR-edited cell lines. F. Cao, E. Ziska, and F.H.U. Gerpott validated the functionality of all cell lines. F. Cao and R. Deliz-Aguirre wrote and validated custom image analysis scripts. F. Cao, and M.J. Taylor performed imaging experiments. F. Cao performed experimental design and data analysis. The project was conceived and supervised by M.J. Taylor. The manuscript was prepared by F. Cao, E. Ziska and M.J. Taylor with input from all authors.

## Materials and Methods

### KEY RESOURCES TABLE

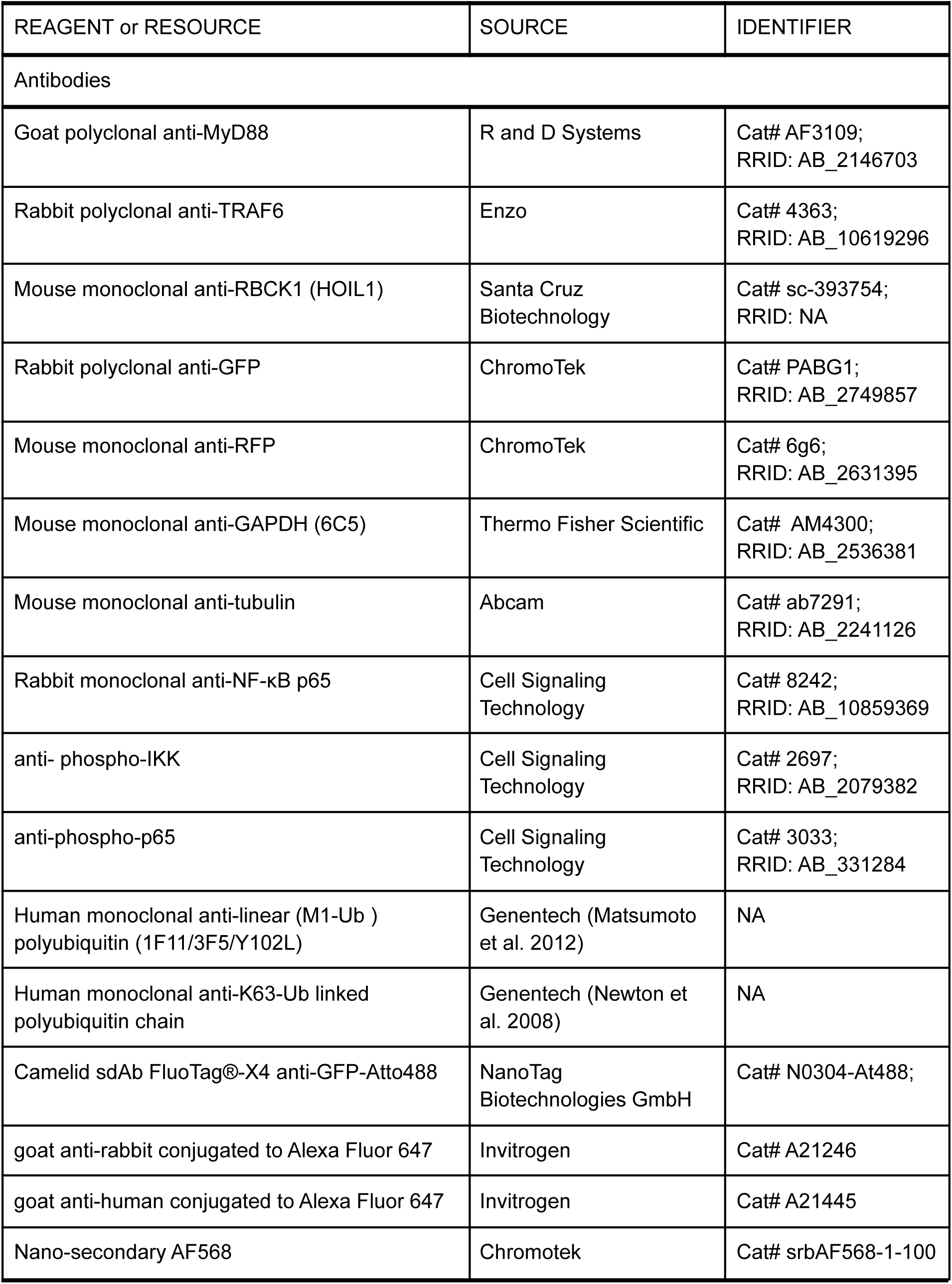
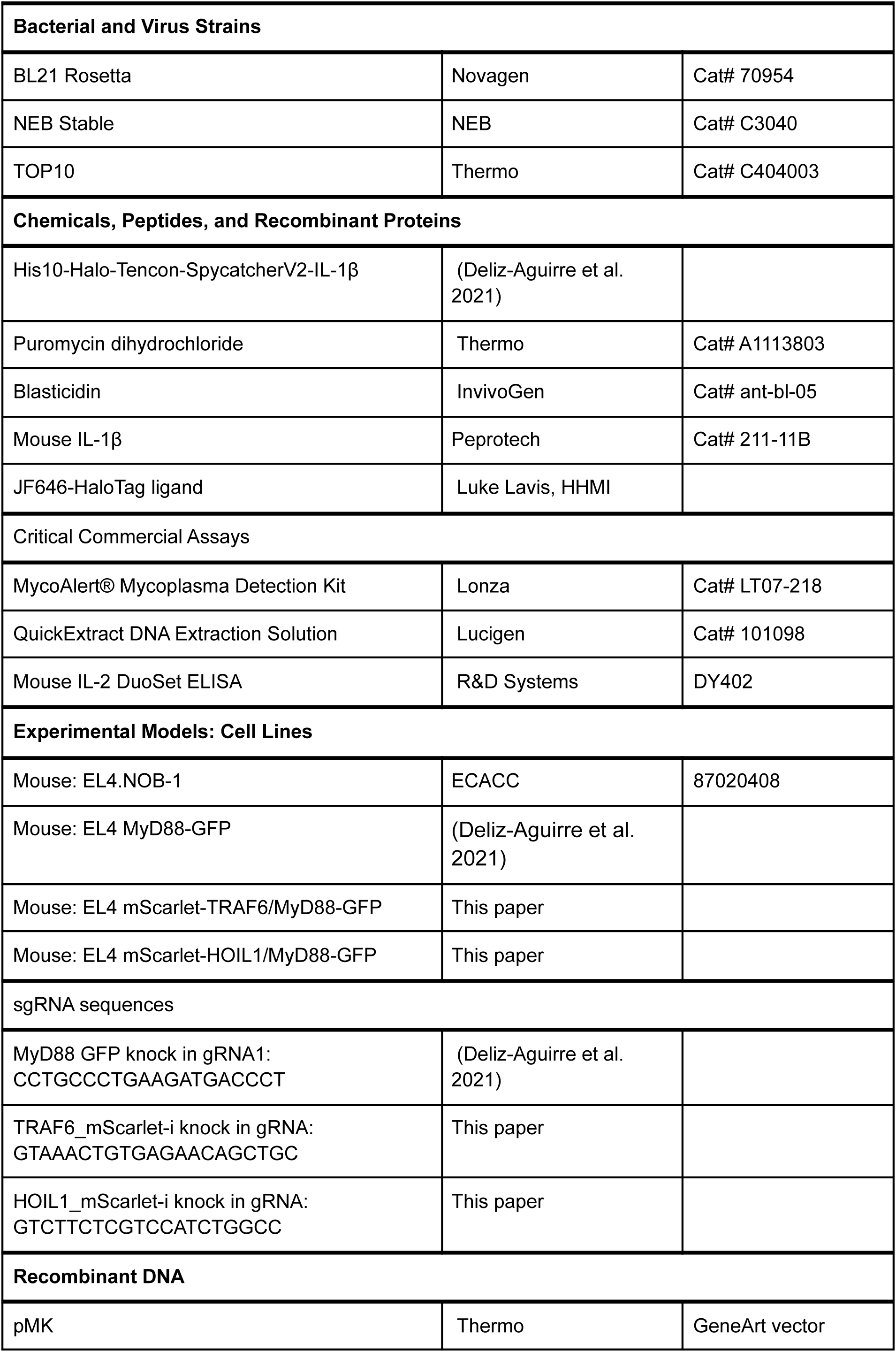
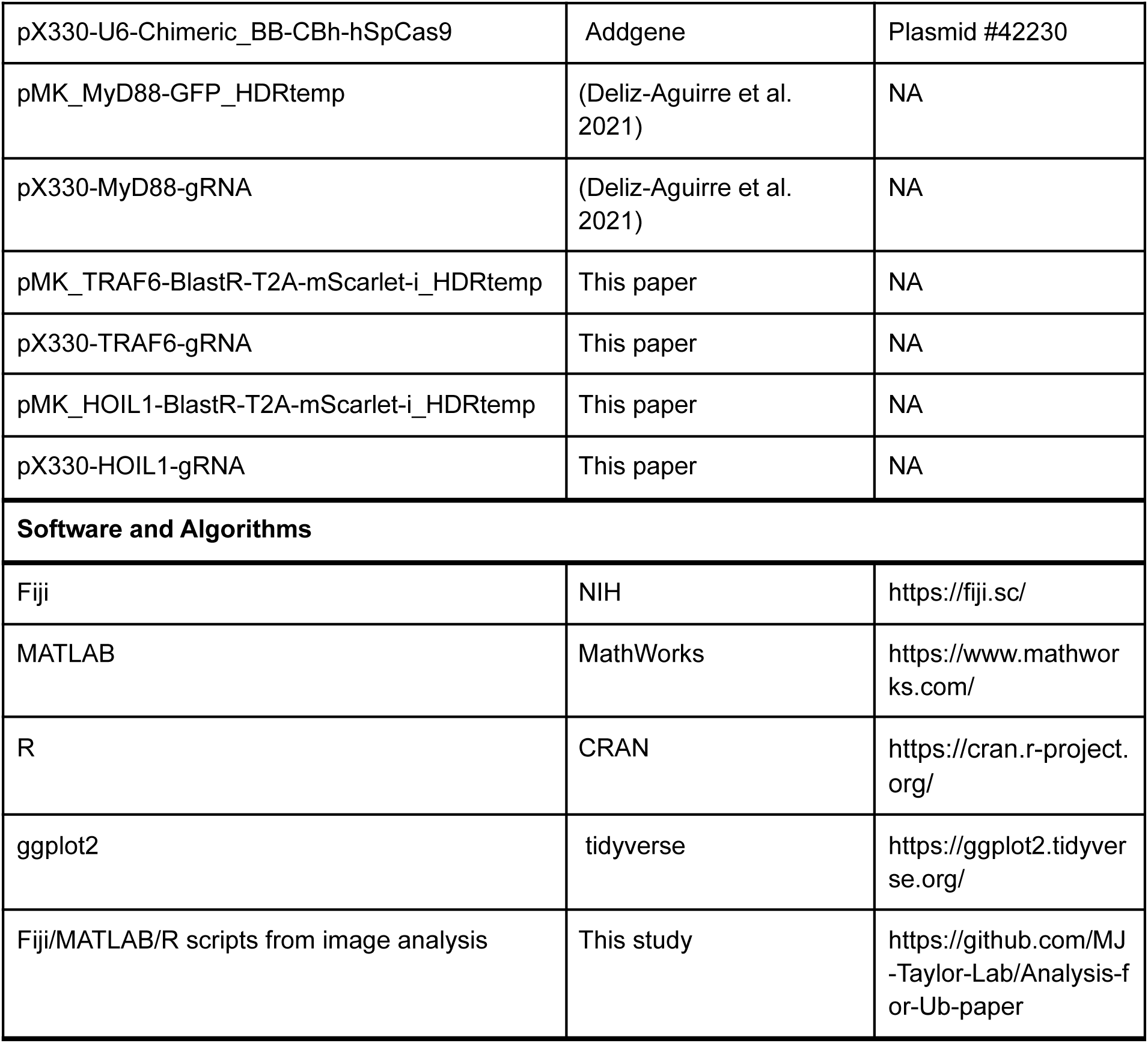

### Methods

#### Contact for Reagent and Resource Sharing

Further information and requests for resources and reagents should be directed to the Lead Contact, Marcus Taylor.

### Experimental Model and Subject Details

#### Cell Culture

EL4.NOB1 WT (purchased from ECACC, and referred to as EL4 in the paper) and gene edited lines (derived from this parental line) were grown in RPMI (Thermo Fisher Scientific) with 10% FBS (Biozol) supplemented with 2 mM L-glutamine. EL4.NOB1 cultures were maintained at a cell density of 0.1-0.5x 10^6^ cells/ml in 5% CO_2_, 37°C. All cells were determined to be negative for mycoplasma (Lonza).

### Method Details

#### Homology-directed repair (HDR) DNA template design for CRISPR/Cas9 **endogenous labeling**

Plasmids DNA repair templates were designed using a pMK (Life Technologies, Carlsbad, CA) vector backbone. Silent mutations were included in the homology arms to remove sgRNA target sites and avoid Cas9 cleavage of the repair template. Homology arms were amplified from EL4 genomic DNA, and assembled with DNA fragments encoding the fluorescent protein tag mScarlet-i (Bindels et al. 2017) and a blasticidin resistance cassette separated by a 2A sequence (Szymczak-Workman, Vignali, and Vignali 2012). These DNA fragments were inserted into a pMK plasmid backbone using Gibson Assembly. All HDR template plasmids were sequence verified. The HDR template to label MyD88 with GFP was described previously (Deliz-Aguirre et al. 2021). Full details of the HDR DNA template plasmid construction used in this study are given below (full sequences of the HDR templates given in Table 1).

#### pMK-BlastR-2A-mScarlet-i-TRAF6-HDR

5’ and 3’ homology arms were designed from the mouse TRAF6 gene (ENSMUSG00000027164) covering a distance of 666 bps and 646 bps either side of the ATG start codon. BlastR-2A-mScarlet-i cassette was inserted between these homology arms and fused to the TRAF6 N-terminus via a 3x Gly-Gly-Ser linker. The following primers were used for the 5’ homology arm:

5’-TTGTAAAACGACGGCCAGCACCAGACTGGGCATTTAGAAATCCACATGGC-3’ and

5’-CTTCTTGAGACAAAGGCTTGGCCATAGTAGCTCTGTTGTCAGTCGATCTTCAg-3’. 3’ homology arm: 5’-GCCAGTCGTCCAGTGACTGCTGCGC-3’ and 5’-GGAAACAGCTATGACCATGCACAAGTAGCTGAGTTTAAGGCAAATTAGAAATACTACTTTC-3’. BlastR-2A-mScarlet-i cassette:

5’-ATGGCCAAGCCTTTGTCTCAAGAAGAAT-3’ and 5’-GCAGCAGTCACTGGACGACTGGCTGGATCCGCAACTGTTCTCACAGTTAAGGAGACTGCTTCCTCCACTGCCTCC-3’. pMK vector backbone:

5’-CATGGTCATAGCTGTTTCCTTGC-3’ and 5’-CTGGCCGTCGTTTTACAACG-3’.

#### pMK-BlastR-2A-mScarlet-i-HOIL1-HDR

5’ and 3’ homology arms were designed from the mouse HOIL1 gene (ENSMUSG00000027466) covering a distance of 598 bps and 790 bps either side of the ATG start codon. A BlastR-2A-mScarlet-i cassette was inserted between these homology arms and fused to the HOIL1 N-terminus via a 3x Gly-Gly-Ser linker. The following primers were used for the 5’ homology arm:

5’-GTTGTAAAACGACGGCCAGCAGAGCTAGGCGCTGCCTGGAGTC-3’ and 5’-ACTGCCTCGCCCTTGCTCACCATCTGGCCTGGCTGCGCCCATCC-3’. 3’ homology arm:

5’-GGCAGTGGAGGAAGCGACGAGAAaACCAAGAAAGGTGGGCACCG-3’ and 5’-GGAAACAGCTATGACCATGTTTTCCGGTCCTCACTCCTCTGTACCC-3’.

BlastR-2A-mScarlet-i cassette:

5’-GGAGGAGAATCCCGGGCCAGTGAGCAAGGGCGAGGCAGTGATC-3’ and 5’-CTCGTCGCTTCCTCCACTGCCTCCTGAGCCACCCTT-3’. pMK vector backbone: 5’-CATGGTCATAGCTGTTTCCTTGC-3’ and 5’-CTGGCCGTCGTTTTACAACG-3’.

#### Generation of CRISPR/Cas9 sgRNA vectors for endogenous labeling of MyD88, **TRAF6 and HOIL1**

Single-guide RNAs (sgRNA) targeting +/-50 bps of the N-terminus start codon of TRAF6 and HOIL1 were designed using the web-based Benchling CRISPR design tool. sgRNAs were selected for each target (see Key Resources Table for sequences), and complementary oligonucleotides designed to be ligated into Bbs1 digested *Streptococcus pyogenes* Cas9 and chimeric guide RNA expression plasmid pX330, (pX330-U6-Chimeric_BB-CBh-hSpCas9, Addgene #42230). sgRNA oligonucleotides were ordered from Integrated DNA Technologies (IDT). Complementary sgRNA oligonucleotides were 5’ phosphorylated with T4 Polynucleotide kinase, annealed, ligated into Bbs1 digested pX330 using Quick Ligase (NEB). pX330 plasmids were transformed into *NEB Stable* competent cells. All sgRNA pX330 plasmids were sequence verified.

#### Generation of CRISPR/Cas9 Engineered Cell Lines

EL4.NOB1 cells were electroporated with a pX330 Cas9/gRNA expressing vector and the pMK vector encoding the HDR template with the Neon Transfection System. EL4 cells were electroporated with the following conditions: voltage (1080 V), width (50 ms), number of pulses (one). For double editing of MyD88/TRAF6 or MyD88/HOIL1 gene loci, 1.5 µg of sgRNA-Cas9 and HDR template plasmids (in equal molar ratio) were electroporated simultaneously. After electroporation, cells were plated in RPMI culture medium without antibiotics for 24 hours. For the selection of TRAF6 and HOIL1 edited alleles, 6 µg/ml blasticidin was added to the cell culture medium 24hr after electroporation. EL4 cells were selected in blasticidin for 48hr.

Monoclonal cell lines were generated by fluorescence-activated cell sorting (FACS). Cells were sorted using BD FACS Aria II at Deutsches Rheuma-Forschungszentrum Berlin, Flow Cytometry Core Facility. To isolate gene-edited EL4 cells, we first performed a bulk sorting of double positive cells. This population was expanded and single cells were sorted into 96-well plates containing culture medium with 15% EL4.NOB-1 conditioned RPMI medium.

The gene edited clonal cell lines were verified using PCR, sequencing and western blot analysis. First, genomic DNA was isolated from selected monoclonal cell lines using QuickExtract DNA Extraction Solution (Epicentre). To test for gene editing and correct insertion of mGFP/mScarlet-i cassette, PCR primers were designed to amplify a DNA fragment that contained the junctions between mGFP/mScarlet-i open reading frame, the 3’ or 5’ homology arm and the gene locus. To check whether single-cell clones were homozygous or heterozygous, we designed PCR primers that amplified a fragment containing mGFP/mScarlet-i cassette, the entire 3’ or 5’ homology arms and the junction between the homology arms and the gene locus (see Table S1). PCR products were analyzed on a 0.8-1% agarose gel, gel extracted using Monarch Nucleic Acid Purification Kits (NEB) and submitted for Sanger Sequencing. Analysis of EL4 HOIL1-mScarlet/MyD88-GFP genomic DNA showed heterozygous editing of the HOIL gene locus and homozygous editing of the MyD88 locus.

To confirm the presence of mEGFP/mScarlet-i fusion protein the cell clones were analyzed by western blot using specific antibodies against MyD88, TRAF6, HOIL1 and GFP or mScarlet-i (RFP). Insertion of fluorescent tags resulted in a 25kDa increase of molecular weight in comparison to non-tagged protein. As expected from the sequencing result, the HOIL1 edited cell line was expressing mScarlet-i-HOIL1 and non-tagged HOIL1. The TRAF6 edited cell line was expressing mScarlet-i-TRAF6 (Fig. S5, full-length western blot shown in Fig. S9). Finally, all cell clones were imaged by microscopy to check for correct localisation of fluorescent signals.

#### Assay of IL-2 release in WT and gene-edited EL4 cells

To measure IL-2 release, we used the Mouse IL-2 DuoSet ELISA kit (R&D Systems; DY402-05) following the manufacturer’s protocol. First 10^6^ cells in 150 μl medium per well were seeded into a 48-well plate and allowed to settle for 30 min. Cells were then stimulated with IL-1β (Peprotech, cat. No. 211-11B) in 50 μl medium per well at a final concentration of 10 ng/10^6^ cells. For unstimulated controls 50 μl medium only was added. After 24 h, plates were centrifuged (300 g for 5 min), and supernatants were transferred to a new plate. Supernatants were stored at −80°C until IL-2-ELISA analysis. Absorbance readings were acquired on a VersaMax Microplate Reader (Molecular Devices) at 450 nm. IL-2 release was assayed on three independent days in triplicate. The obtained results were normalized based on the EL4 WT IL-2 release (Fig. S5C).

#### Chromium nanopatterned coverslips

Chromium nanopatterned coverslips with the design and specification described (see Fig. S1) were produced by ThunderNIL Srl (Trieste, Italy). Coverslips were fabricated by the pulsed nanoimprint lithography method (W.-C. Lin et al. 2010) and printed with a master design which contained multiple nanopatterned chromium grids containing square corrals with 2.5 or 1 µm^2^ dimensions. Chromium gridlines were 100 nm thick and 5 nm high and were printed on no.1.5 coverslips with a diameter of 25 mm.

#### Imaging Chambers and Supported Lipid Bilayers

SLBs were prepared using a previously published method (Taylor et al. 2017; Deliz-Aguirre et al. 2021). Briefly, Phospholipid mixtures consisting of 97.5% mol 1-palmitoyl-2-oleoyl-*sn*-glycero-3-phosphocholine (POPC), 2% mol 1,2-dioleoyl-sn-glycero-3-[(N-(5-amino-1-carboxypentyl)iminodiacetic acid)succinyl] (ammonium salt) (DGS-NTA) and 0.5% mol 1,2-dioleoyl-sn-glycero-3-phosphoethanolamine-N-[methoxy(polyethylene glycol)-5000] (PE-PEG5000) were mixed in glass round-bottom flasks and dried down with a rotary evaporator. All lipids used were purchased from Avanti Polar Lipids. Dried lipids were placed under vacuum for 2 hrs to remove trace chloroform and resuspended in PBS. Small unilamellar vesicles (SUVs) were produced by several freeze-thaw cycles combined with bath sonication. Once the suspension had cleared, the SUVs were spun in a benchtop ultracentrifuge at 35,000xg for 45 min. SUVs were stored at 4°C for up to a week. IL-1β-functionalized SLBs were formed in 96-well glass bottom plates (Matrical), coverslips (25 mm diameter, No. 1.5 H, Marienfeld-Superior) or on nanopatterned coverslips (25 mm diameter, No. 1.5 H, grid lines produced by ThunderNIL Srl).

To prepare SLBs on 96-well glass bottom plates (Matrical), the plates were cleaned for 30 min with a 5% Hellmanex solution containing 10% isopropanol heated to 50°C, then incubated with 5% Hellmanex solution for 1 h at 50°C, followed by extensive washing with pure water. 96-well plates were dried with nitrogen gas and sealed until needed. To prepare SLB, individual wells were cut out and base etched for 15 min with 5 M KOH and then washed with PBS. To form SLBs, SUVs suspension was deposited in each well or coverslip and allowed to form for 1 hr at 45°C. After 1 hr, wells were washed extensively with PBS. SLBs were incubated for 15 min with HEPES buffered saline (HBS: 20 mM HEPES, 135 mM NaCl, 4 mM KCl, 10 mM glucose, 1 mM CaCl_2_, 0.5 mM MgCl_2_) with 10 mM NiCl_2_ to charge the DGS-NTA lipid with nickel. The SLBs were then washed in HBS containing 0.1% BSA to block the surface and minimize non-specific protein adsorption. After blocking, the SLBs were functionalized by incubation for 1 hr with His10-IL-1β. The labeling solution was then washed out and each well was completely filled with HBS with 0.1% BSA. For SLBs set up on 96-well plates the total well volume was 630 μl (manufacturers specifications), and 530 μl was removed leaving 100 μl of HBS 0.1% BSA in each well. Each SLBs was functionalized with 100 µl His10-Halo-IL-1β of twofold desired concentration for 1 hr and excessive ligands were washed away with HBS.

To prepare SLBs on normal or nanopatterned coverslips, the coverslips were cleaned by bath-sonication for 30 min in MilliQ H2O. After sonication, coverslips were immersed in freshly prepared piranha solution (sulfuric acid:hydrogen peroxide, 3:1) for 15 min, rinsed in MilliQ water 20 times and finally dried with nitrogen gas. To form SLBs, we sandwiched 30 µl of a SUVs suspension between a petri dish and a coverslip. After a 5 min incubation the petri dish was immersed in MilliQ water bath. The coverslip was removed from the petri dish, and washed in the MilliQ water to remove excessive SUVs. Coverslips were assembled in Attofluor Chamber (Thermo Fisher). The MilliQ water in each chamber was slowly replaced with PBS and incubated with 10 mM NiCl_2_ for 15 min, followed by incubation with 0.1% BSA for 30 min. Finally, each SLBs was functionalized with His10-Halo-IL-1β for 1 hr and excessive ligands were washed away with 20 ml HBS.

#### Protein Expression, Purification and Labeling

To functionalize the SLBs with active mouse IL-1β, we expressed and purified fusion protein of His10-Halo-IL-1β as previously described (Deliz-Aguirre et al. 2021).

This protein was produced from two separate expression plasmids:

pET28a-MmIL1β-Spytag and pET28a-His10-Halo-Tencon-SpycatcherV2. We expressed IL-1β-Spytag and His10-Halo-Tencon-SpycatcherV2 in BL21-DE3 Rosetta *E. coli* (Novagen) grown in Terrific Broth media. After an overnight induction with IPTG, the bacterial culture was pelleted and the cell pellets was resuspended in the lysis buffer (50 mM TRIS pH 8.0, 250 mM NaCl, 5 mM Imidazole with protease inhibitors, Lysozyme 100 µg/ml) and lysed using sonication. To covalently couple His10-Halo-Tencon-Spycatcher to MmIL1β-Spytag, the cleared lysates were mixed and incubated with mild agitation for 1 hour at 4 C. To ensure complete spycatcher-spytag conjugation, the lysates were mixed with 2:1 ratio (vol:vol, based on starting bacterial culture volume) of MmIL1β-Spytag to His10-Halo-Tencon-Spycatcher. After the conjugation, the His10-Halo-Tencon-Spycatcher-IL-1β-Spytag was purified by Ni-NTA resin. Conjugation was monitored by mobility shift using SDS-PAGE. After elution, the protein was desalted with HiTrap desalting column into 20 mM HEPES and subject to anion exchange chromatography with a MonoQ column. This was followed by gel filtration over Superdex 200 26/600 into storage buffer (20 mM HEPES, 150 mM NaCl). The protein was snap frozen with the addition of 20% glycerol in liquid nitrogen and placed at −80°C for long-term storage. In text this protein is referred to as His10-Halo-IL-1β. Following purification, the His10-Halo-Tencon-Spycatcher-IL-1β-spytag protein was either snap-frozen and stored at –80°C or directly used for HaloTag-labeling. To label the HaloTag, a 2.5x molar excess of JF646-HaloLigand was mixed with the protein and incubated at room temperature for 1 hour followed by an overnight incubation at 4°C. Post labeling, the protein was gel filtered over a Superdex 200 26/600 into storage buffer and snap-frozen with the addition of 20% glycerol in liquid nitrogen and placed in −80°C for storage. The degree of labeling was calculated with a spectrophotometer by comparing 280 nm and 640 nm absorbance (usually 85-95% labeling efficiency was achieved).

For microscopy calibration of mScarlet single molecule intensity we used His10-mScarlet-IL-1β (previously described here(Deliz-Aguirre et al. 2021)). For mEGFP single molecule intensity, His10-mEGFP was expressed from a pET28a vector, and purified with Ni-NTA resin followed by gel filtration. Frozen aliquot of both proteins were stored at -80C.

#### Immunofluorescence staining and Widefield microscopy of RelA nuclear translocation

To analyze the nuclear translocation of RelA in IL-1β-stimulated EL4 cells with or without inhibition of Myddosome coalescence (Fig. 1G), IL-1β-functionalized SLBs were prepared on coverslips without chromium grid lines (off grid) or on coverslips with 2.5 µm or 1 µm grid lines. Non-functionalized SLBs served as unstimulated controls. EL4 were incubated for 30 min with IL-1β-labeled SLBs functionalized with 100 IL-1β molecules/µm^2^. Cells were then fixed with 3.5% (wt/vol) PFA containing 0.5% (wt/vol) Triton X-100 for 20 min at room temperature. Cells were washed with PBS and blocked with PBS 10% BSA (wt/vol) at 4°C overnight.

The next day, fixed cells were labeled with a one-step staining solution containing anti-RelA (1:400, Cell Signaling Technology, #8242), Nano-secondary AF568 (Chromotek, #srbAF568-1-100, 1:1000) diluted in PBS 10% (wt/vol) BSA containing 0.1% Triton X-100, for 1 h at room temperature. Cells were labeled with DAPI and 488-phalloidin to label the nucleus and cytosol respectively. Finally, cells were washed five times in PBS before imaging.

We acquired Widefield microscopy images of RelA nuclear translocation on an inverted microscope (Nikon TiE) equipped with Lumencor Spectra-X illumination. Fluorescent images were acquired with Nikon Plan Apo 40x 0.95 NA air objective lens and projected onto a Photometric Prime 95 camera and a 1.5x magnification lens (calculated pixel size of 181.41 nm). Image acquisition was performed with NIS-Elements software.

#### Immunofluorescence staining of phospho-p65, phospho-IKK, K63-Ub and M1-Ub

To analyze the colocalization of phospho-IKK, K63-Ub and M1-Ub with MyD88-GFP (Fig. 2), EL4 cells were stimulated with IL-1β-functionalized SLBs for 30 mins, and then fixed with 3.5% (wt/vol) PFA containing 0.5% (wt/vol) Triton X-100 for 20 min at room temperature. Staining was then performed with a traditional two-step staining method. After fixation, cells were washed with PBS, then blocked in PBS 10% (wt/vol) BSA containing 4% normal goat serum for 1 h at room temperature. Fixed cells were labeled with primary antibodies diluted in PBS 10% (wt/vol) BSA containing 0.1% Triton X-100 at 4 C overnight. The next day, cells were washed five times with PBS and labeled with secondary antibodies (goat anti-rabbit/human conjugated to Alexa Fluor 647; 1:1000; Invitrogen, #A21246/A21445) and FluoTag-X4 anti-GFP conjugated to Atto488 (1:500, Nano Tag Biotechnology, #N0304-At488-L) for 1 hr at room temperature. Finally, cells were washed five times in PBS before imaging with TIRF or SIM microscopy. To analyze the colocalization of phospho-p65 with MyD88 we used a one-step staining protocol detailed below. To image pIKK with SIM (Fig. 2C), coverslips were mounted in prolong glass antifade mountant (Thermo, #P36980).

To quantitatively compare phospho-p65, phospho-IKK, K63-Ub and M1-Ub staining off and on grids (Fig. 3-4), EL4-MyD88-GFP cells were starved in serum-free medium for 3-4 hr. After starvation, 7x10^5^ cells were applied to SLBs and incubated at 37°C for 30 min. We used SLB functionalized with 5-25 IL-1β/µm^2^. Cells were fixed with a final concentration of 3.5% (wt/vol) PFA containing 0.5% (wt/vol) Triton X-100 for 20 min at room temperature. Cells were washed with PBS and blocked with PBS 10% BSA (wt/vol) at 4°C overnight. The next day, we performed one-step staining. Cells were incubated with a antibody mixture containing primary antibody, Nano-secondary AF568 (Chromotek, #srbAF568-1-100, 1:1000) prepared in PBS with 10% (wt/vol) BSA and 0.1% Triton X-100. Cells were incubated with this labeling solution at room temperature for 1 hr. Across all immunofluorescence experiments we used the following primary antibodies and labeling concentrations: anti-phospho-p65, 1:800 (Cell Signaling Technology, #3033); anti-phospho-IKK, 1:400 (Cell Signaling Technology, #2697); anti-M1-Ub, 1:2500 ((Matsumoto et al. 2012), Genentech); anti-K63, 1:10000 ((Newton et al. 2008), Genentech). Cells were then labeled with FluoTag-X4 anti-GFP conjugated to Atto488 at room temperature for 1 hr. After antibody labeling each chamber was washed with 40 ml PBS. Cells were then imaged immediately with TIRF microscopy.

#### TIRF-Microscopy data acquisition

Imaging of MyD88-GFP, mScarlet-TRAF6, and mScarlet-HOIL1 recruitment was performed on an inverted microscope (Nikon TiE) equipped with a Nikon fiber launch TIRF illuminator. Illumination was controlled with a laser combiner using the 488-, 561-, and 640-nm laser lines at ∼0.35, ∼0.25, and ∼0.17 mW laser power, respectively (laser power measured after the objective). Fluorescence emission was collected through filters for GFP (525 ± 25 nm), RFP (595 ± 25 nm), and JF646 (700 ± 75 nm). All images were collected using a Nikon Plan Apo 100x 1.4 NA oil immersion objective that projected onto a Photometrics 95B Prime sCMOS camera with 2 x 2 binning (calculated pixel size of 150 nm) and a 1.5x magnifying lens. Image acquisition was performed using NIS-Elements software. All experiments were performed at 37°C. The microscope stage temperature was maintained using an OKO Labs heated microscope enclosure. Images were acquired with an interval of 4 s using exposure times of 60–100 ms.

#### Imaging EL4 cells endogenously expressing MyD88-GFP, mScarlet-TRAF6 or **mScarlet-HOIL1 on IL-1β functionalized SLBs with TIRF-Microscopy**

His10-Halo-JF646-IL-1β-functionalized SLBs were set up as described above. To quantify the density of IL-1β on the SLB, wells were prepared that were functionalized with identical labeling protein concentration and time, but with different molar ratios of labeled to unlabeled His10-Halo-IL-1β. Before application of cells, SLBs were analyzed by TIRF microscopy to check formation, mobility, and uniformity. Short time series were collected at wells containing a ratio of labeled to unlabeled His10-HaloIL-1β (e.g., <1 His10-Halo-JF646-IL-1β molecule/µm^2^) to calculate ligand densities on the SLB based upon direct single molecule counting. By controlling the concentration of His10-Halo-JF646-IL-1β in the labeling reaction we could label SLB with final IL-1β densities ranging from 1-200 molecules/µm^2^.

Before each imaging experiment, we acquired calibration images using recombinant mEGFP and His10-mScarlet-IL-1β previously described here (Deliz-Aguirre et al. 2021). To image a single GFP/mScarlet-i fluorophores, the recombinant purified proteins were diluted in HBS and adsorbed to KOH-cleaned glass. Single molecules of GFP/mScarlet-i were imaged using identical microscope acquisition settings to those used for cellular imaging. To image live cells, EL4 cells were pipetted onto supported lipids bilayers functionalized with His10-Halo-JF646-IL-1β. EL4 cells expressing MyD88-GFP, mScarlet-TRAF6, or mScarlet-HOIL1 were sequentially illuminated for 60-100 ms with 488-nm and 561-nm laser line at a frame interval of 4 s (Fig. 5). Diffraction-limited punctate structures of MyD88-GFP, mScarlet-TRAF6, or mScarlet-HOIL1 were detected and tracked using the Fiji TrackMate plugin (Tinevez et al. 2017).

#### Structured Illumination Microscopy data acquisition

We acquired 3D structured illumination microscopy of fixed EL4 cells on a Zeiss Elyra 7 microscope equipped with 405, 488, 561 and 642 nm laser lines for excitation. Image acquisition was performed with a 63x, NA 1.46 oil objective and images were captured on pco.edge 4.2 sCMOS camera. We acquired Z stacks of fixed cells using the 3D leap acquisition plugin using a 200 nm Z axis step size with 13 phases. We performed post-processing image reconstruction in Zeiss Zen software.

#### Quantification and Statistical Analysis

All data are expressed as the mean ± the standard deviation (SD) or mean ± the standard error of the mean (SEM), as stated in the figure legends and results. The exact value of n and what n represents (e.g., number of cells, MyD88-GFP puncta, or experimental replicates) is stated in figure legends and results. Means of experimental replicates were compared using an unpaired two-tailed Student’s t-test implemented in R studio. Data distribution was assumed to be normal based on density plots, but this was not formally tested. The scripts used in this study for image analysis are available at https://github.com/MJ-Taylor-Lab/Analysis-for-Ub-paper. All data used in this study is available at 10.5281/zenodo.7408704.

### Quantification of immunofluorescence staining of RelA nuclear localization

We quantified widefield microscopy images of RelA nuclear localization in an analysis pipeline implemented in FIJI and Cell Profiler. First we performed background subtraction from the MyD88-GFP and RelA (Cy5 channel) immunofluorescence staining micrographs in FIJI. Background was removed in two steps: First we subtracted a dark field image from each image. Secondly we estimate cytosolic background by generating median blur from each micrograph. We then subtracted this median blur from the parent micrograph.

We then performed segmentation and quantification using a custom Cell Profiler pipeline that allowed images to be processed in batch. We segmented the cell nucleus using the DAPI channel. Selected nuclei retained for analysis had to have a diameter between 30 to 60 pixels, this excluded small DAPI stained objects that corresponded to cell fragments and apoptotic cells. Segmentation of the 488-phalloidin staining channel identified the total cell volume. Both segmentation steps were performed using an Otsu threshold. The volume corresponding to cellular cytoplasm is identified by subtracted the total cell volume minus nucleus volume. The RelA staining intensity of the cell nucleus and cytoplasm was extracted, and the ratio calculated. RelA nucleus to cytoplasm ratio from images acquired on 2.5 µm and 1 µm grids and unstimulated negative controls were normalized to the intensity of RelA nucleus to cytoplasm ratio from off grids data. We normalized intensity using the following equation: *Norm. Int = (Intensity - quantile(0.05)_off grid_)/(quantile(0.95)_off grid_ - quantile(0.05)_off grid_)*. Finally, we performed data visualization of normalized RelA nucleus to cytoplasm ratio in ggplot2 (Fig. 1H).

### Quantification of immunofluorescence staining and analysis of phospho-p65, phospho-IKK, K63-Ub and M1-Ub

We quantified TIRF microscopy images of pp65, pIKK, K63-Ub and M1-Ub immunofluorescence staining in an analysis pipeline implemented in FIJI and Cellprofiler (McQuin et al. 2018). First we performed background subtraction from the MyB88-GFP and immunofluorescence staining TIRF micrographs in FIJI. Background was removed in two steps: First we subtracted a dark field image from each image. Secondly we estimate cytosolic background by generating median blur from each TIRF micrograph. We then subtracted this median blur from the parent TIRF micrograph.

Next we segmented MyD88-GFP puncta and quantified fluorescence intensity using a custom Cell Profiler pipeline that allowed images to be processed in batch. We segmented MyD88-GFP puncta using Otsu threshold. Only segmented MyD88-GFP puncta were retained that has a diameter between 3 to 30 pixels. After image segmentation and object detection the integrated intensity and mean intensity of the MyD88-GFP and immunofluorescence staining channel were extracted for each segmented puncta. We performed manual inspection of the segmented images and objects to verify correct processing and remove incorrectly segmented puncta.

Data normalization and visualization were performed using R. To compare MyD88-GFP puncta size and staining intensity across different replicates acquired on different days we normalized puncta fluorescence intensities. Fluorescence intensities of MyD88-GFP and immunofluorescence staining from images acquired on 2.5 µm and 1 µm grids were normalized to the intensity of those from off grids data. We normalized intensity using the following equation: *Norm. Int = (Intensity - quantile(0.01)_off grid_)/(quantile(0.99)_off grid_ - quantile(0.01)_off grid_)*.

We used the following criteria to classify MyD88 puncta in fixed cells as single Myddosomes or clusters of Myddosomes (Fig, 3G, H and 4G, H). We observed that MyD88 puncta that formed on grids rarely had a fluorescent intensity greater than 0.5 (normalized integrated intensity, Fig. 3E, F). In contrast we found that off grid, between 3-5% of MyD88 puncta were classified as clusters (Fig. S3C, D and 4C, D). This was in agreement with our live cell measurement of MyD88 puncta size (Fig. 1F). Based on these observations and previous measurement that nanopatterned grids disrupted cluster formation (Fig. 1F), we defined MyD88 puncta that were clusters as puncta with an intensity ≥0.5, and single Myddosomes as puncta being below this threshold. Finally, we performed data visualization of MyD88-GFP puncta size and immunofluorescence staining intensity in ggplot2 (Fig. 3A-B, E-H and 4A-B, E-H).

### Quantification and analysis of MyD88-GFP puncta and colocalization and recruitment of mScarlet-TRAF6/mScarlet-HOIL1

To quantify the dynamics of MyD88-GFP, mScarlet-TRAF6 and mScarlet-HOIL1, we used an image analysis pipeline described previously (Deliz-Aguirre et al. 2021) and briefly described here. First, images in each channel were processed in Fiji to remove background fluorescence. Background subtraction was performed in two steps. First, we subtracted a dark frame image (acquired with no light exposure to the camera, but identical exposure time to experimental acquisition) to remove noise intrinsic to the camera. Then we subtracted a median-filtered image (generated in Fiji from a median blurred image generated with a radius of 25 pixels) to remove the background associated with cytosolic fluorescence. Next, individual cells were segmented in Fiji according to a maximum projection of MyD88-GFP fluorescence channel. After segmentation, we tracked MyD88-GFP and mSclarlet-TRAF6/mScarlet-HOIL1 puncta in each cell using the Fiji Trackmate plugin (Tinevez et al. 2017).

Tracking coordinates generated by Trackmate were imported into MATLAB, and the fluorescence intensity of MyD88-GFP puncta was measured from a 3x3 pixel region. To quantify colocalization between MyD88-GFP and mScarlet-TRAF6/HOIL1 puncta, we used the tracking coordinates to identify puncta that colocalized for at least two or more consecutive frames. Colocalized puncta were defined as having centroids ≤0.25 µm apart at a given time point. By these criteria, MyD88 tracked puncta were classified as either positive or negative for TRAF6/HOIL1 colocalization.

To estimate the size and number of MyD88s in MyD88-GFP puncta, we acquired images of single mEGFP fluorophores (referred to as simply GFP) absorbed to glass with identical imaging settings to those used in live cell imaging. Images of single molecules of mEGFP were processed identically to live cell imaging data, with background subtraction, tracking and intensity measurement performed as described above. Once intensity measurements were obtained for single molecules of GFP, this was used to divide the fluorescence intensity of MyD88-GFP puncta to yield an estimate of MyD88 copy number. To normalize the puncta by the number of Myddosome complexes, we divided GFP normalized intensity by 4.5 (e.g., the intensity of a single Myddosomse, based on the broad fluorescent intensity distribution of a complex containing 6x MyD88-GFP (S.-C. Lin, Lo, and Wu 2010), see Fig. S6B, and (Deliz-Aguirre et al. 2021)). Using this criteria a MyD88 puncta is defined as a Myddosome complex if the fluorescent intensity is greater than or equal to 4.5x GFP, and is a cluster of Myddosomes (e.g., 2 or more complex) if the fluorescent intensity is greater than or equal to 9x mean intensity of GFP.

Finally, we performed data analysis and visualization in R. MyD88-GFP puncta with an intensity ≥4.5x the mean intensity of mEGFP were defined as fully assembled Myddosome complexes (see (Deliz-Aguirre et al. 2021)). Myddosome clusters (defined as MyD88-GFP puncta containing 2 or more Myddosome complexes), were defined as MyD88-GFP puncta with an intensity ≥9x the mean intensity of mEGFP (see Fig. S6B).

### Quantification of mScarlet-TRAF6/HOIL1 lifetime

To quantify mScarlet-TRAF6/HOIL1 lifetime, the max number of Myddosome clusters was rounded to integers and was binned into three different categories (single Myddosomes, 2-4 Myddosomes and 5-7 Myddosomes). The lifetime of TRAF6 and HOIL1 puncta that colocalized with those specific Myddosome clusters was plotted as histograms (Fig 7A). We measured 1845 and 2302 lifetimes for TRAF6 recruitment to single Myddosomes, 1427 and 713 lifetimes at clusters composed of 2-4 Myddosomes, 91 and 26 lifetimes at clusters composed of 5-7 Myddosomes, off and on grids, respectively (10.4% and 8.4% of total single Myddosomes, 21.2% and 25.7% of total 2-4 Myddosomes, and 28.8% and 38.8% of total 5-7 Myddosomes off and on grids at 10 IL-1/µm^2^). We measured 446 and 575 lifetimes of HOIL1 recruitment to single Myddosomes, 2051 and 1171 lifetimes at clusters composed of 2-4 Myddosomes, and 1130 and 275 lifetimes at clusters composed of 5-7 Myddosomes, off and on grids, respectively (1% and 1.8% of total single Myddosomes, 6.4% and 13.3% of total 2-4 Myddosomes, and 11.5% and 40.4% of total 5-7 Myddosomes off and on grids at 32 IL-1/µm^2^).

## Supplementary Movies Movie 1

**Movie 1.**
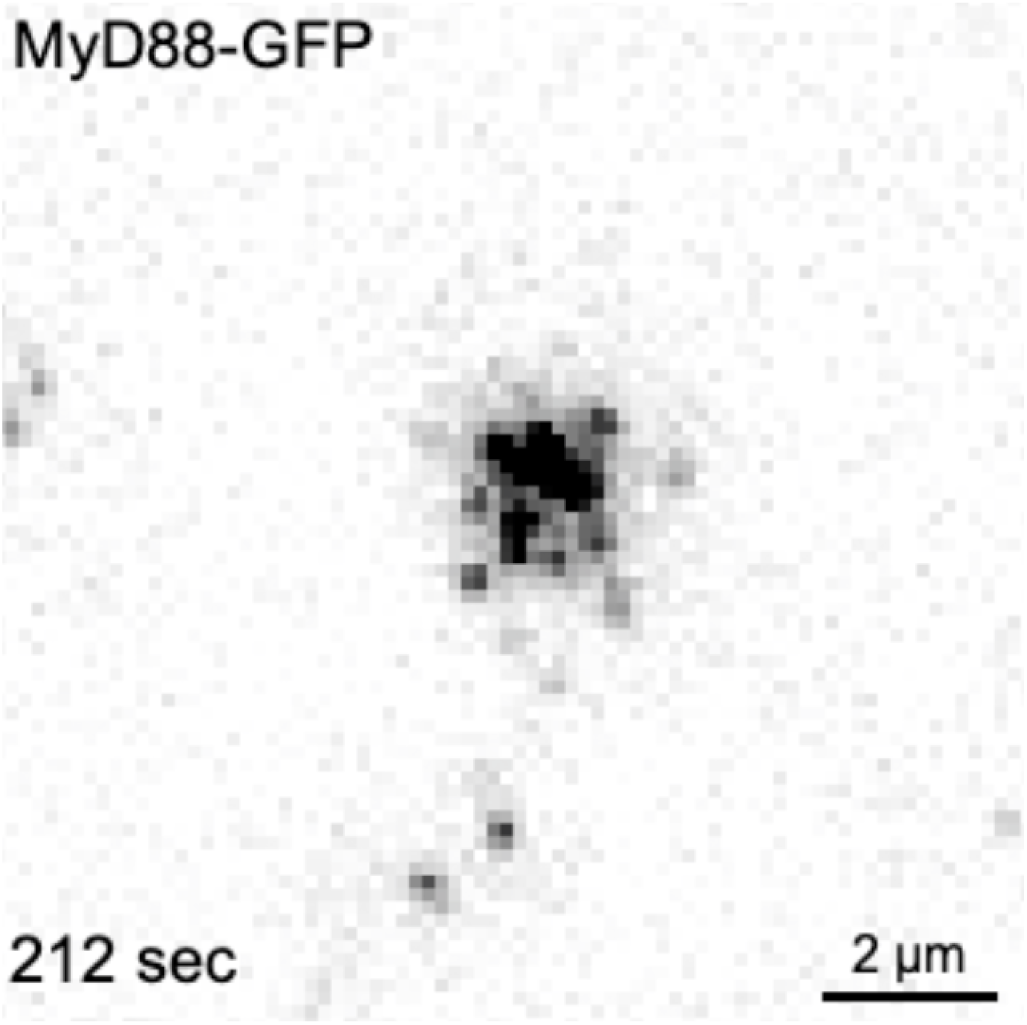
Myddosome clustering on IL-1-functionalized supported lipid bilayers (relates to **Fig. 1A**). This movie shows an EL4-MyD88-GFP cell interacting with IL-1-functionalized supported lipid bilayers. This movie illustrates that MyD88-GFP is recruited to the plasma membrane upon stimulation. MyD88-GFP assembles into dynamic puncta that coalesce into large clusters at the cell:SLB interface.

**Movie 2.**
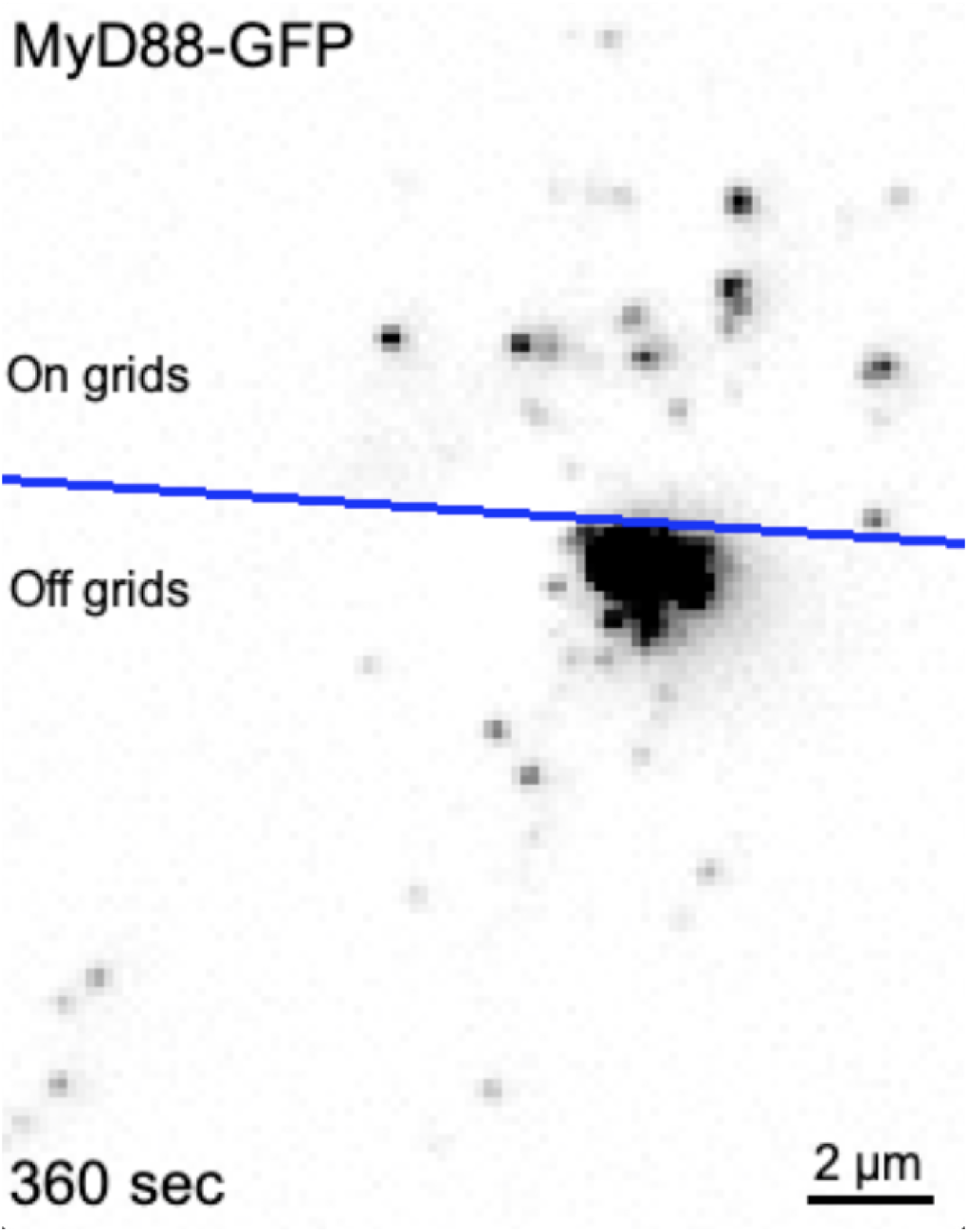
Myddosome clustering with or without the presence of extracellular physical barriers (relates to **Fig. 1D**). This movie shows an EL4-MyD88-GFP cell landing at the boundary between a continuous SLB (off grids) and a partitioned SLB formed on a nano-patterned chromium grid. The boundary between the continuous and gridded SLB is marked by the blue line on the movie. MyD88-GFP puncta off grid coalesce to form a large cluster. However, the mobility of MyD88-GFP puncta on grids is restricted to individual corrals and these puncta are unable to merge with MyD88-GFP puncta in adjacent corrals. This movie illustrates that Myddosomes are tethered to the cell surface and extracellular barriers that restrict the mobility of SLB tethered IL-1, can in turn restrict the coalescence of intracellular Myddosomes.

**Movie 3:**
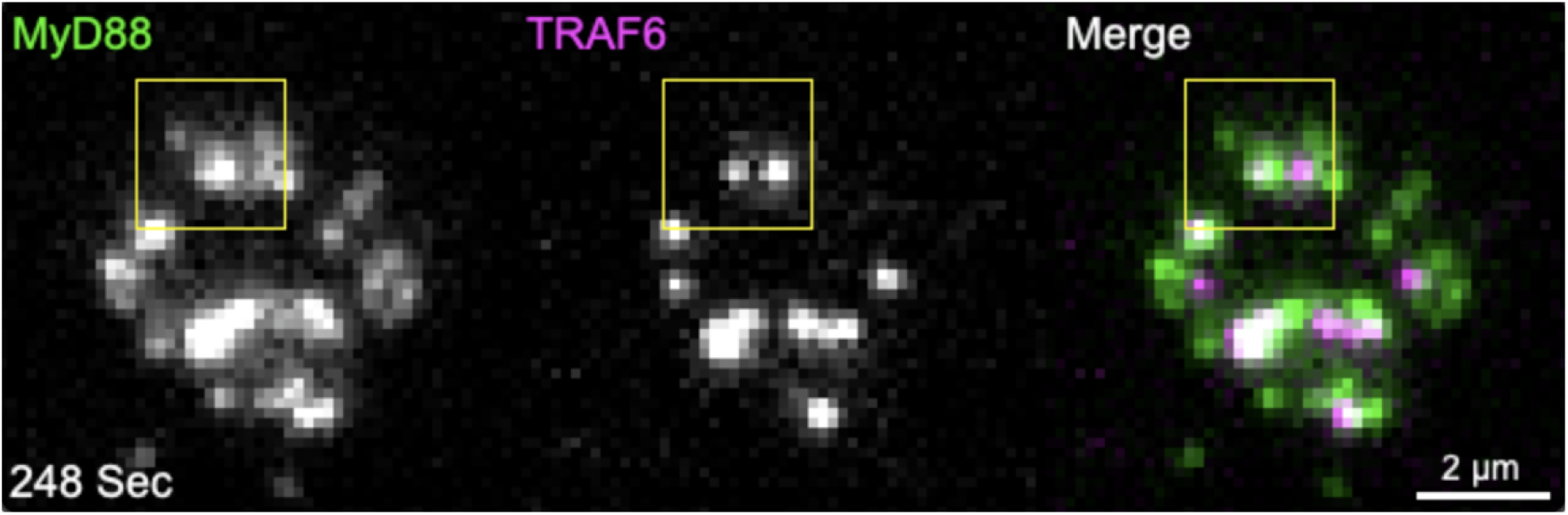
Recruitment of E3 ligase TRAF6 (relates to **Fig. 5A**). This movie shows an EL4 cell endogenously expressing MyD88-GFP and mScarlet-TRAF6 interacting with IL-1 functionalized SLBs imaged with TIRF microscopy. Yellow square region of interest corresponds to the colocalized puncta of MyD88 and TRAF6 shown in the montage in Fig. 5A. This movie illustrates that mScarlet-TRAF6 is recruited to Myddosomes on the plasma membrane.

**Movie 4:**
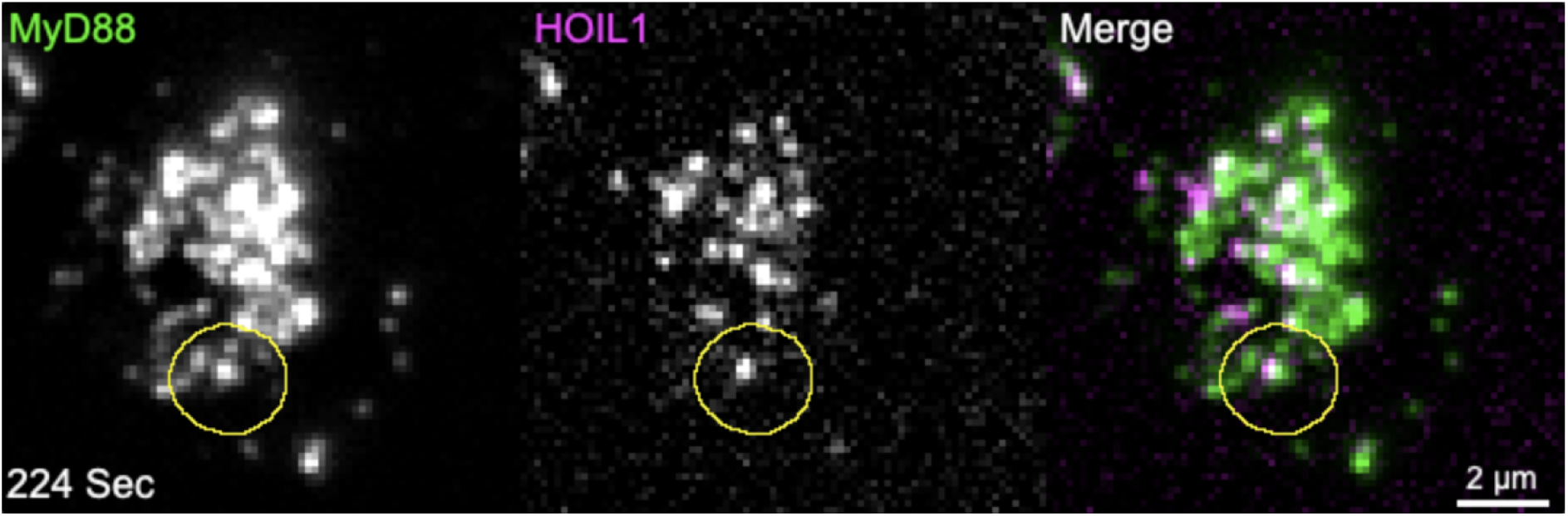
Recruitment of E3 ligase HOIL1 (relates to **Fig. 5B**). This movie shows an EL4 cell endogenously expressing MyD88-GFP and mScarlet-HOIL1 interacting with IL-1 functionalized SLBs and imaged with TIRF microscopy. Yellow circle region of interest is centered on the colocalized puncta of MyD88 and HOIL1 shown in the montage in Fig. 5B. This movie illustrates that mScarlet-HOIL1 is recruited to the Myddosome on the plasma membrane.

**Movie 5:**
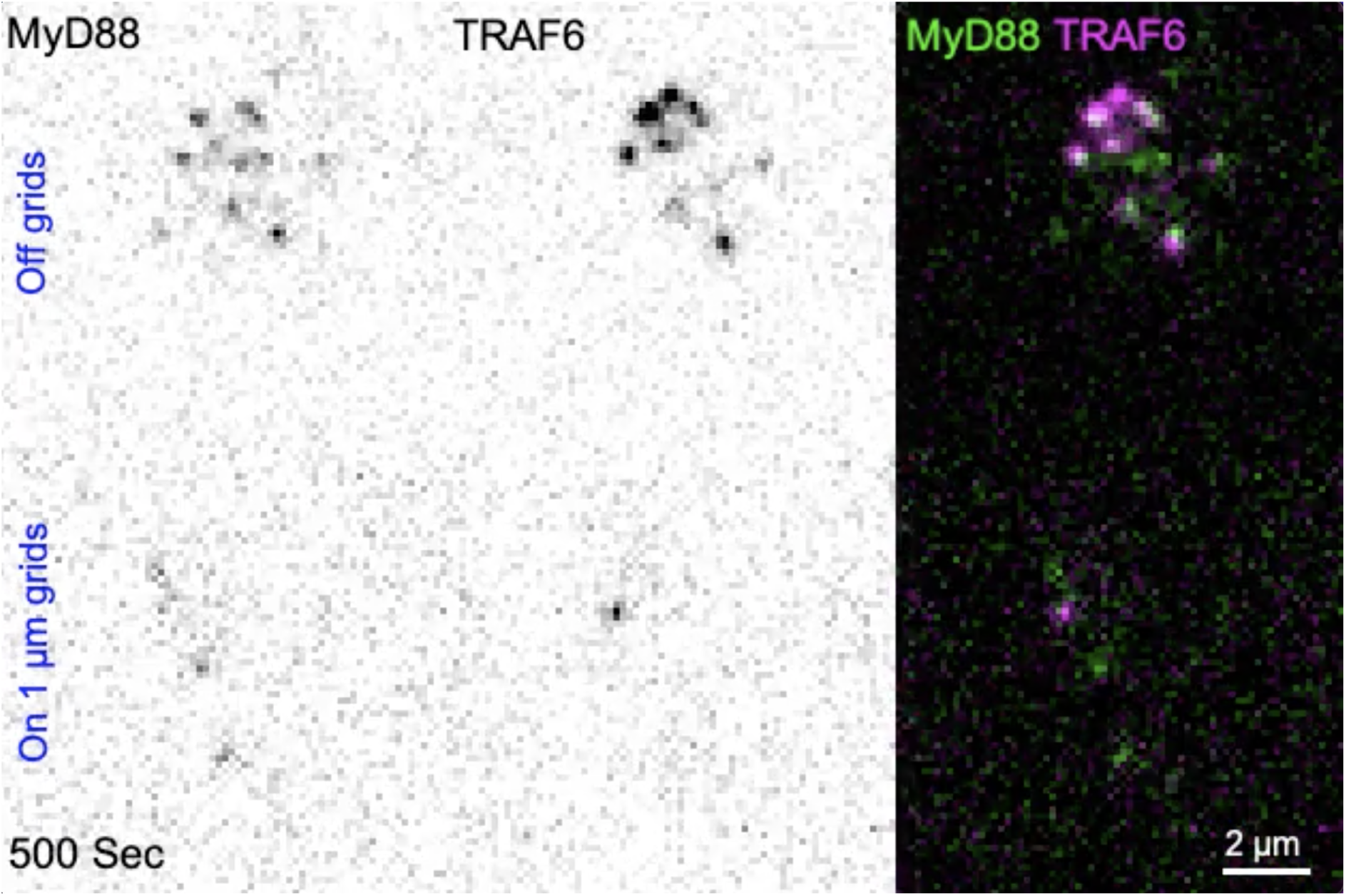
TRAF6 recruitment off and on 1 µm grids (relates to **Fig. 6A** and **6B**). This movie shows EL4 cells endogenously expressing MyD88-GFP and mScarlet-TRAF6 interacting with continuous IL-1-functionalized SLBs (top movie) and 1 µm grid partitioned SLBs (bottom movie) at a density of 1 IL-1/µm^2^. The movie was acquired using TIRF microscopy.

**Movie 6:**
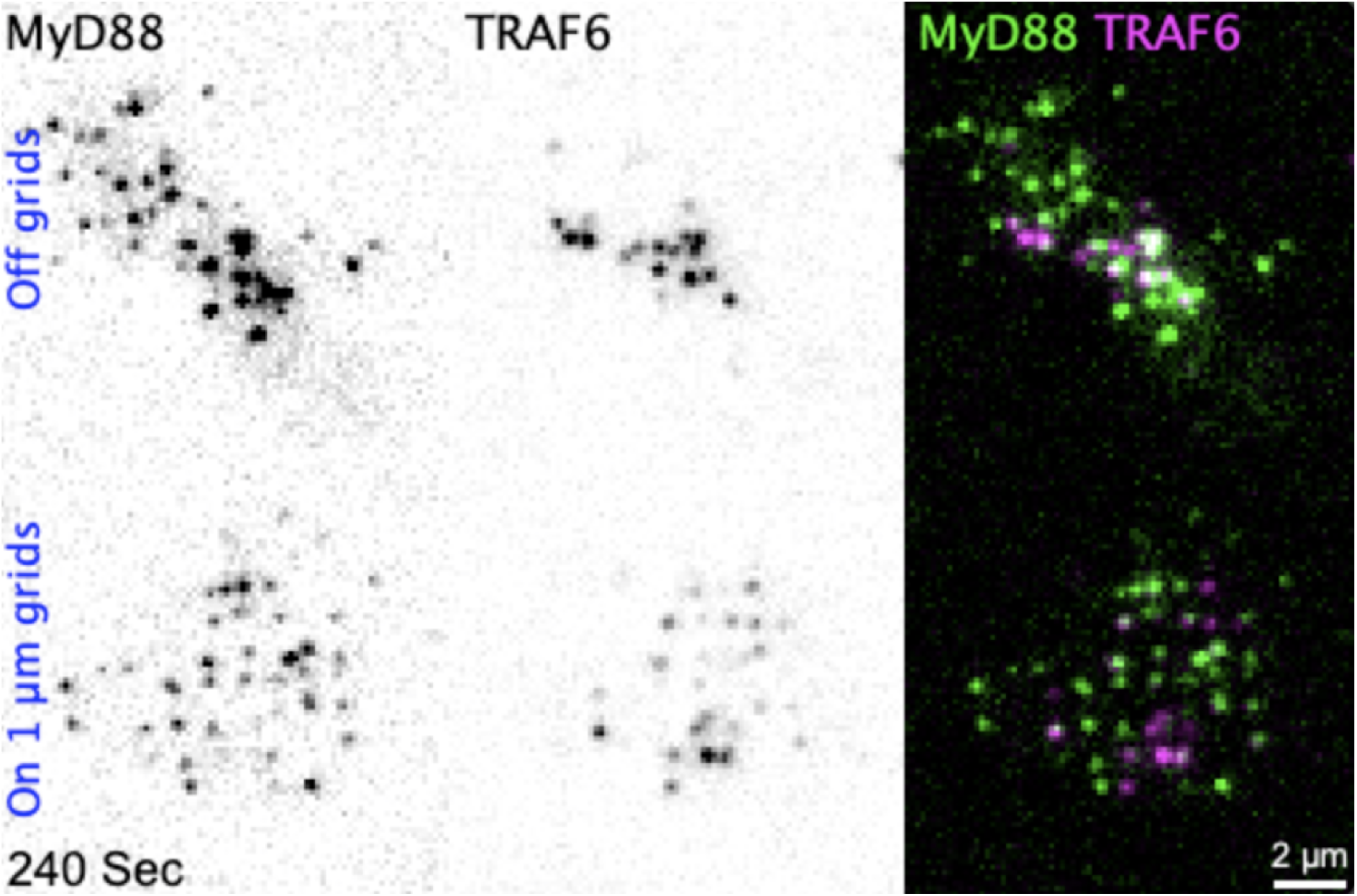
TRAF6 recruitment off and on 1 µm grids (relates to **Fig. 6D** and **6E**). This movie shows EL4 cells endogenously expressing MyD88-GFP and mScarlet-TRAF6 interacting with continuous IL-1-functionalized SLBs (top movie) and partitioned SLBs (bottom movie) at a density of 10 IL-1/µm^2^. The movie was acquired using TIRF microscopy.

**Movie 7:**
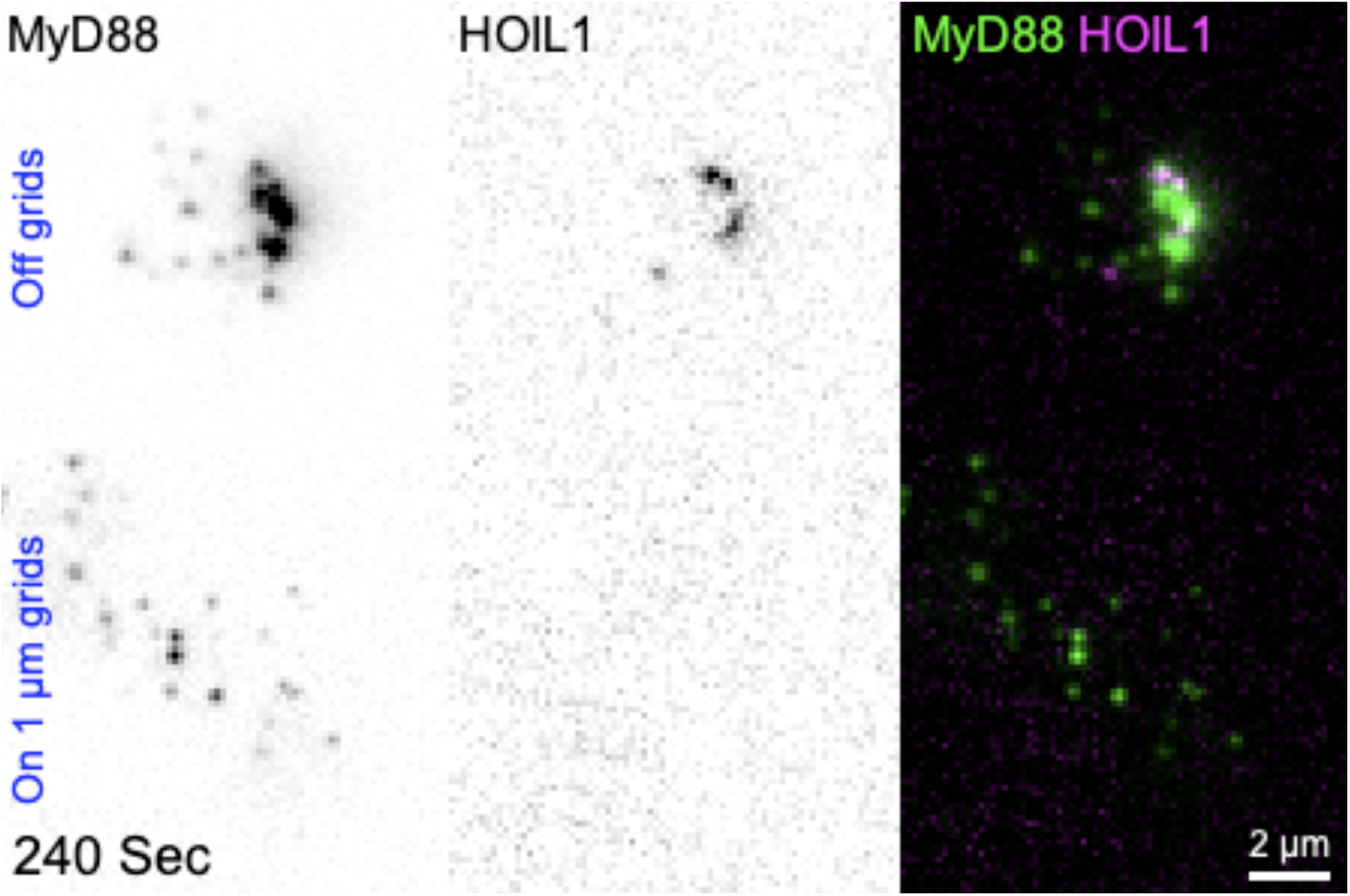
HOIL1 recruitment off and on 1 µm grids (relates to **Fig. 6G** and **6H**). This movie shows EL4 cells endogenously expressing MyD88-GFP and mScarlet-HOIL1 interacting with continuous IL-1-functionalized SLBs (top movie) and partitioned SLBs (bottom movie) at a density of 10 IL-1/µm^2^. This movie illustrates that 1 µm grids diminish the recruitment of mScarlet-HOIL1 to Myddosomes. The movie was acquired using TIRF microscopy.

**Movie 8:**
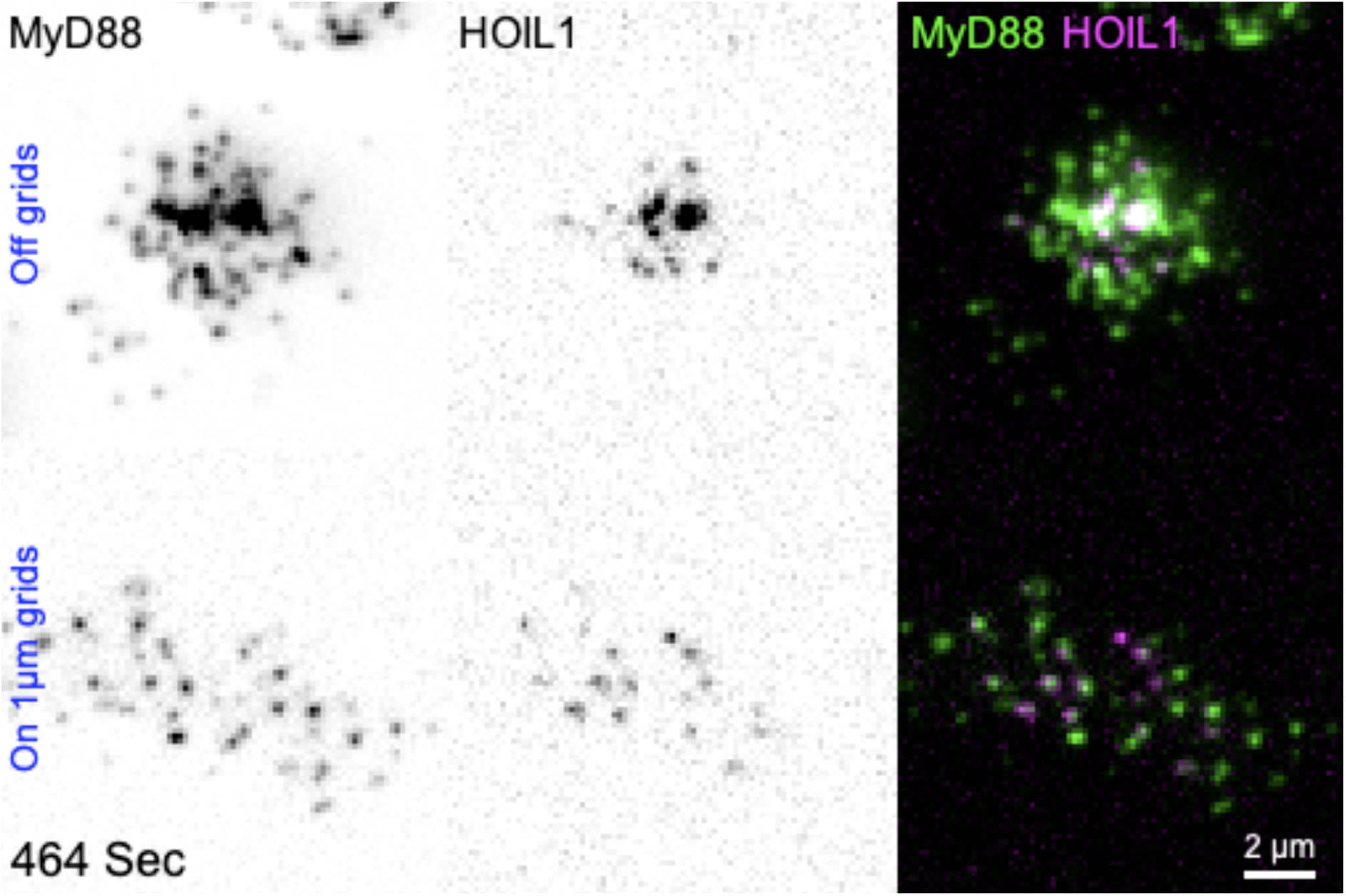
HOIL1 recruitment off and on 1 µm grids (relates to **Fig. 6J** and **6K**). This movie shows EL4 cells endogenously expressing MyD88-GFP and mScarlet-HOIL1 interacting with continuous IL-1-functionalized SLBs and partitioned SLBs at a density of 32 IL-1/µm^2^. This movie illustrates that increasing IL1 density can rescue the recruitment of mScarlet-HOIL1 to MyD88-GFP on 1 µm grids. The movie was acquired using TIRF microscopy.

**Figure S1.**
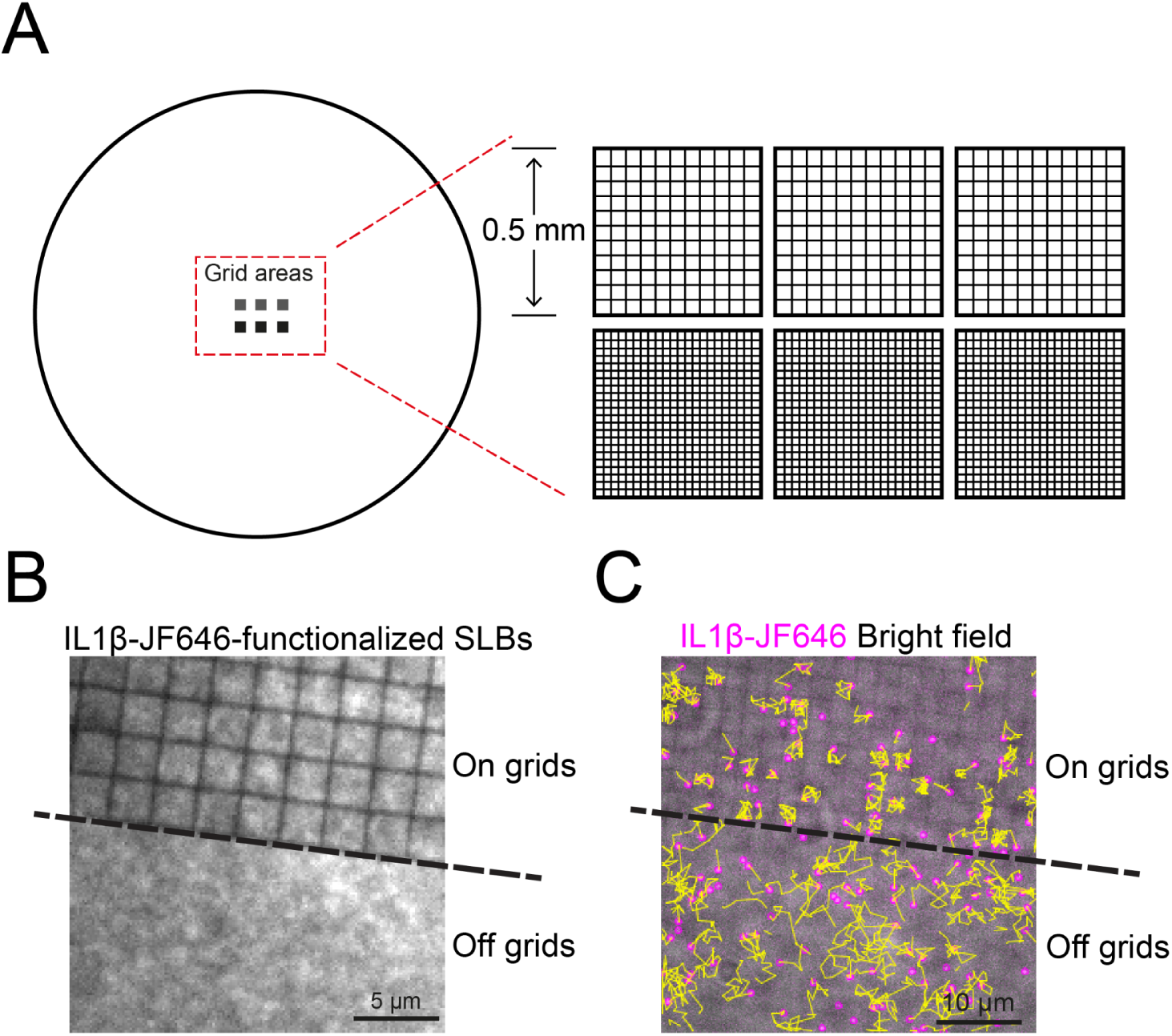
Illustrations of nanopatterned coverslips to assemble partitioned SLBs. (A) Schematic of the coverslip used for preparing partitioned SLBs. The no. 1.5 coverslips with a diameter of 25 mm were fabricated with nanopatterned chromium grids containing square corrals with 2.5 or 1 µm^2^ dimensions. Each dimension contains 3 pieces of square grids with a side length of 0.5 mm. (B) TIRF image showing IL-1-JF646 functionalized SLBs at a field of view containing on grids and off grids SLBs. Continuous SLBs are observed off grids and on each square corral on grids. (C) Mobility of IL-1 ligands off and on grids. Yellow lines are trajectories of IL-1-JF646. Off grids, IL-1 can freely move around, while on grids, IL-1 can only move within individual corrals.

**Figure S2.**
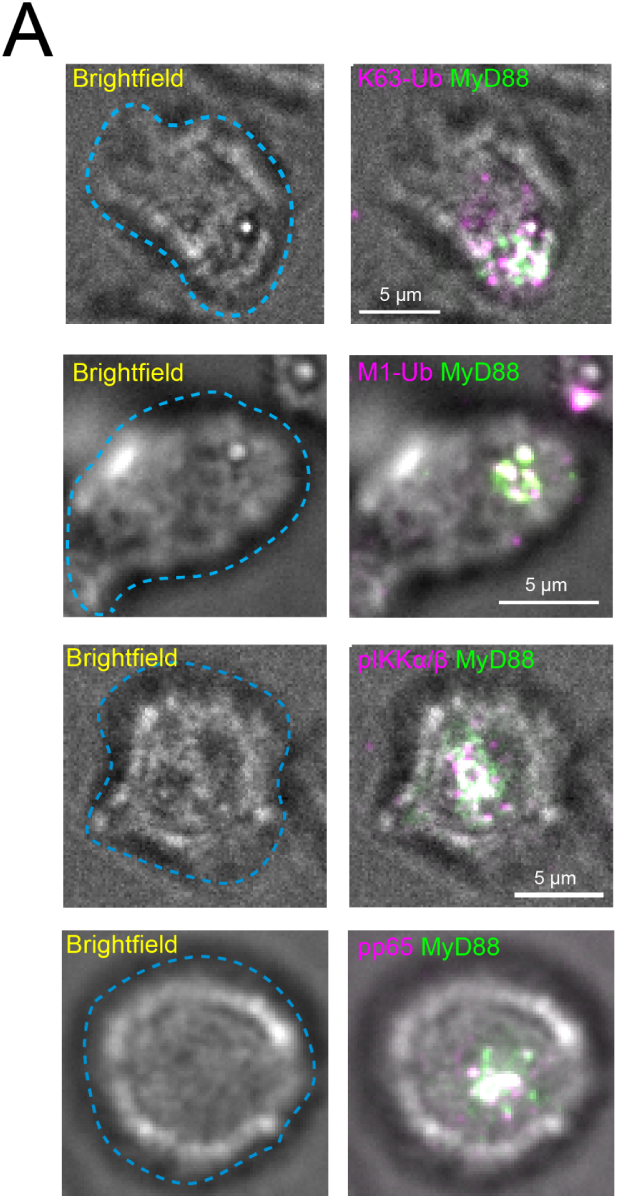
Brightfield images from **Fig. 2A**. Left row are brightfield images overlaid with cell contour (blue dashed line). Right row are brightfield images overlaid with MyD88-GFP and antibody staining.

**Figure S3.**
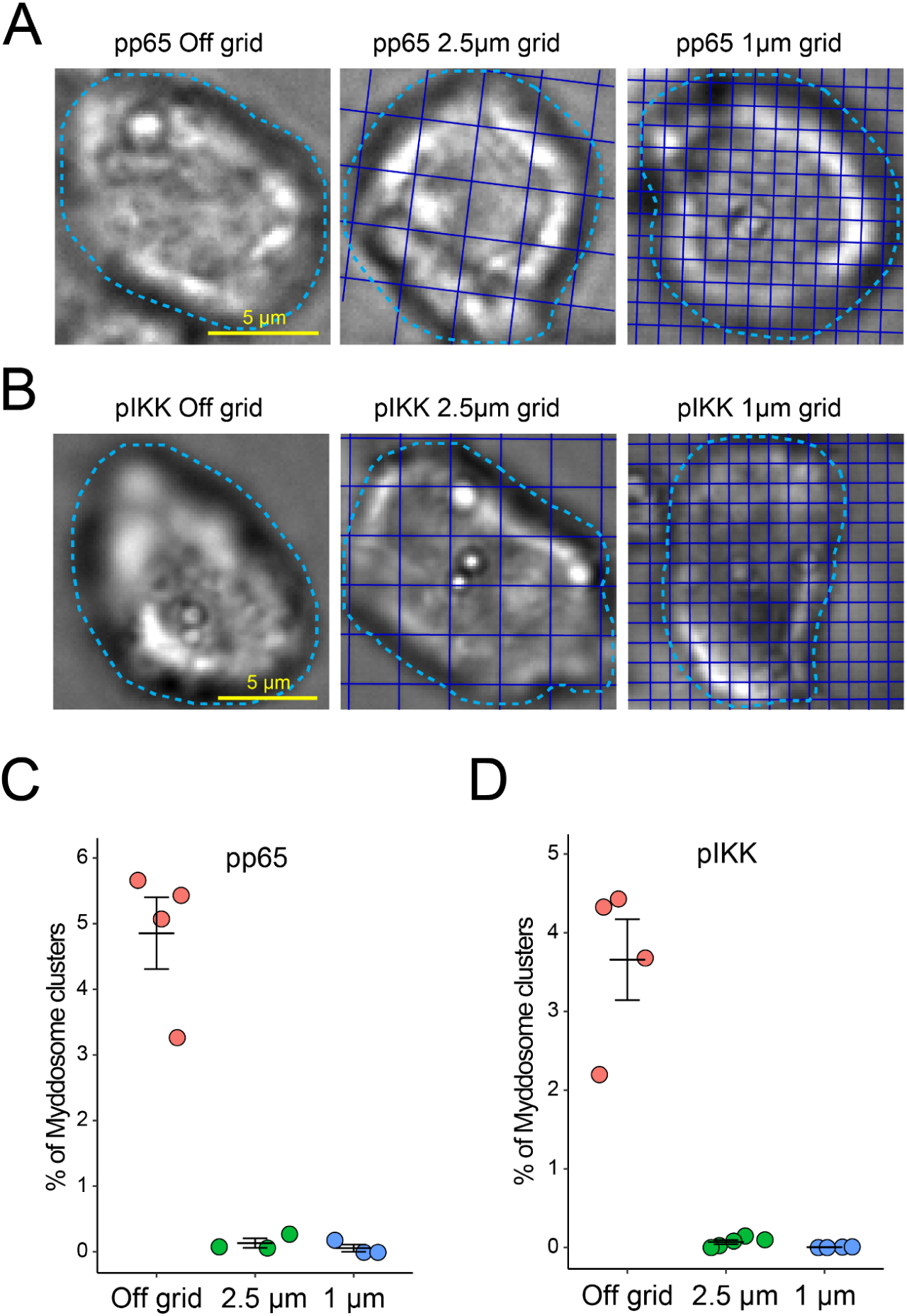
Quantification of pp65 and pIKK staining. (A and B) Brightfield images for Fig. 3A and 3C (A), and Fig. 3B and 3D (B). Cyan dashed lines are cell contour and blue lines indicate grid lines. (C and D) Quantification of the percentage of MyD88 puncta classified as Myddosome clusters in pp65 (C) or pIKK (D) immunofluorescence staining off grids, on 2.5 µm and on 1 µm grids. MyD88 puncta with an integrated intensity greater than or equal to 0.5 are classified as Myddosome clusters. The percentage of Myddosome clusters off grids, on 2.5 µm and on 1 µm grids in pp65 staining are 4.853 ± 0.547%, 0.133 ± 0.072% and 0.055 ± 0.055%, respectively; and that in pIKK staining are 3.657 ± 0.515%, 0.071 ± 0.026% and 0.004 ± 0.003%, respectively. Bars represent mean ± SEM (n = 3 - 4 experimental replicates for pp65, with 259 - 9342 puncta measured per replicate, and 10273, 14009 and 2675 puncta measured off grids, on 2.5 µm and 1 µm grids, respectively; n = 4 - 5 experimental replicates for pIKK, with 435 - 24998 puncta measured per replicate, and 2375, 35496 and 59593 puncta measured off grids, on 2.5 µm and 1 µm grids, respectively)

**Figure S4.**
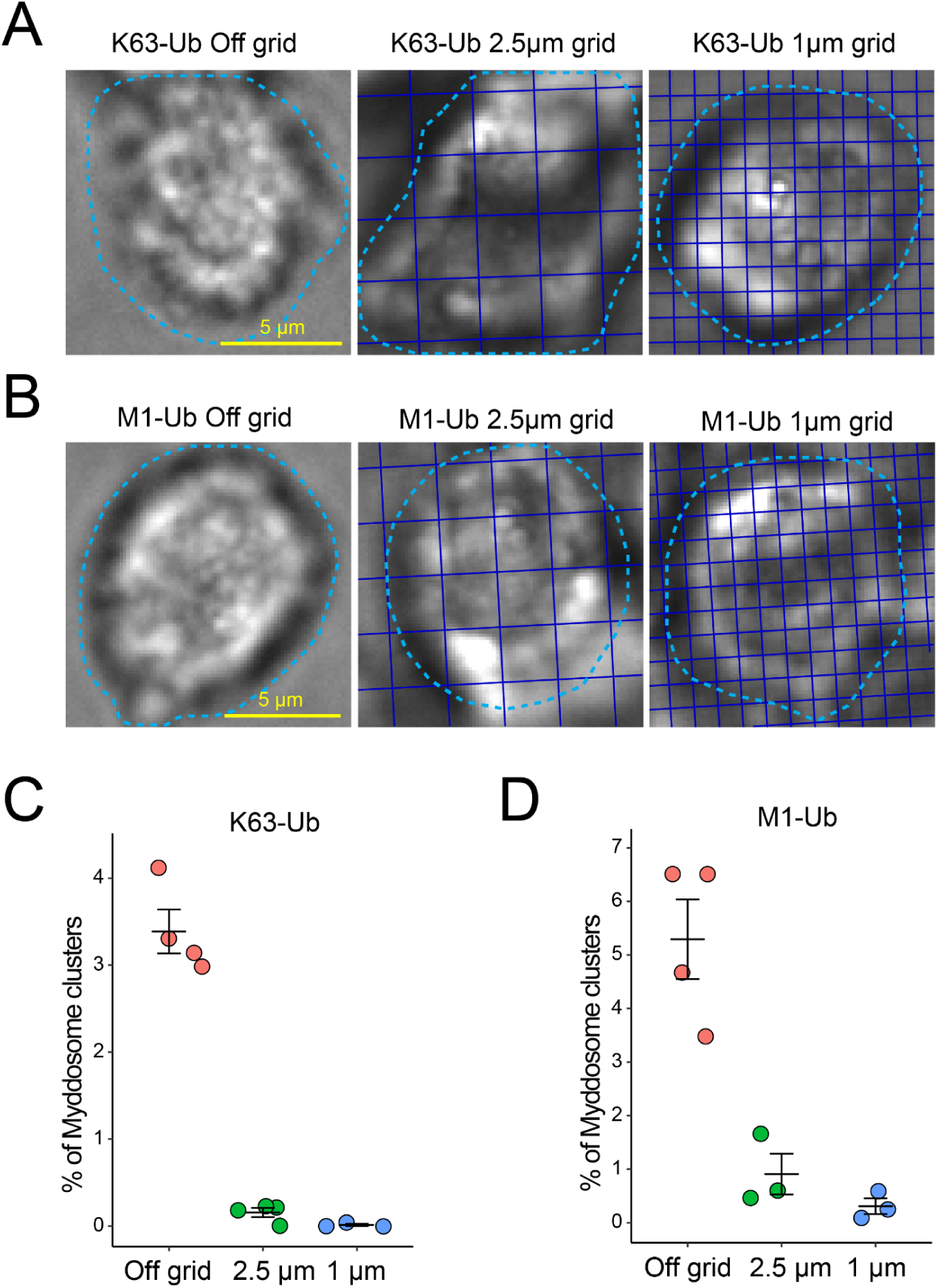
Quantification of K63-Ub and M1-Ub staining. (A and B) Brightfield images for Fig. 4A and 4C (A), and Fig. 4B and 4D (B). Cyan dashed lines are cell contour and blue lines indicate grid lines. (C and D) Quantification of the percentage of MyD88 puncta classified as Myddosome clusters in K63-Ub (C) or M1-Ub (D) immunofluorescence staining off grids, on 2.5 µm and on 1 µm grids. MyD88 assemblies with an integrated intensity greater than or equal to 0.5 are classified as large MyD88 assemblies. The percentage of large MyD88 assemblies off grids, on 2.5 µm and on 1 µm grids in K63-Ub staining are 3.387 ± 0.253%, 0.156 ± 0.053% and 0.013 ± 0.013%, respectively; and that in M1-Ub staining are 5.292 ± 0.744%, 0.908 ± 0.378% and 0.310 ± 0.147%, respectively. Bars represent mean ± SEM (n = 3 - 4 experimental replicates for K63-Ub, with 484 - 14628 puncta measured per replicate, and 14571, 27494 and 24026 puncta measured off grids, on 2.5 µm and 1 µm grids, respectively; n = 3 - 4 experimental replicates for M1-Ub, with 338 - 3887 puncta measured per replicate, and 3114, 6091 and 1844 puncta measured off grids, on 2.5 µm and 1 µm grids, respectively)

**Figure S5.**
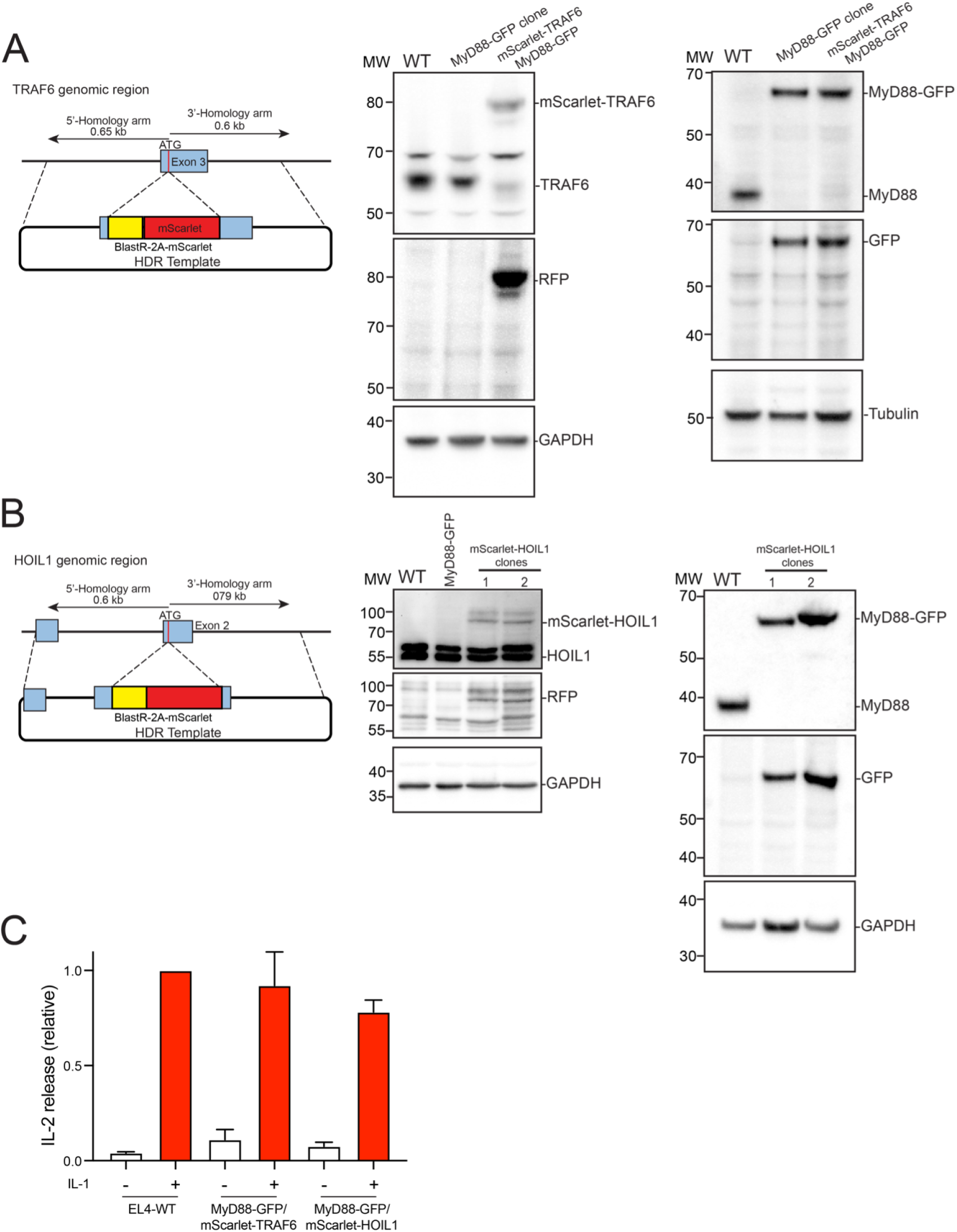
Validation of EL4 cell lines with CRISPR/Cas9 gene edited MyD88-GFP and mScarlet-TRAF6/HOIL1. (A and B) Left, schematic of the TRAF6 (A) or HOIL1 (B) gene locus and HDR template designed to insert a mScarlet open reading frame immediately after the start codon. EL4 cells were electroporated with HDR and gRNA/Cas9 plasmids to simultaneously edit MyD88 and TRAF6 (A) or MyD88 and HOIL1 (B) gene loci. Middle, western blot analysis of mScarlet-TRAF6 (A) or HOIL1 (B) in dual gene-edited EL4 clones. Western blots of whole cell lysates were probed with anti-TRAF6/HOIL1 and anti-RFP antibodies to confirm editing and insertion of fluorescent protein open reading frames at both gene loci. WT EL4 cells and EL4 cells expressing MyD88-GFP only served as negative controls. Right, western blot analysis of MyD88-GFP in dual gene-edited EL4 clones. Western blots of same lysates were probed with anti-MyD88 and anti-GFP to confirm editing and insertion of GFP open reading frame at the MyD88 gene locus. WT EL4 cells served as a negative control and EL4 cells expressing MyD88-GFP only served as a positive control. For MyD88-GFP/mScarlet-HOIL1 cells, all data presented were acquired with clone 1. (C) IL-2 release in WT and dual gene-edited EL4 cells. IL-2 release was measured by ELISA 24 h after IL-1β stimulation. Values for gene-edited cells shown relative to EL4 WT. Average values are calculated from 3-4 independent experiments. Bars represent mean ± SEM.

**Figure S6.**
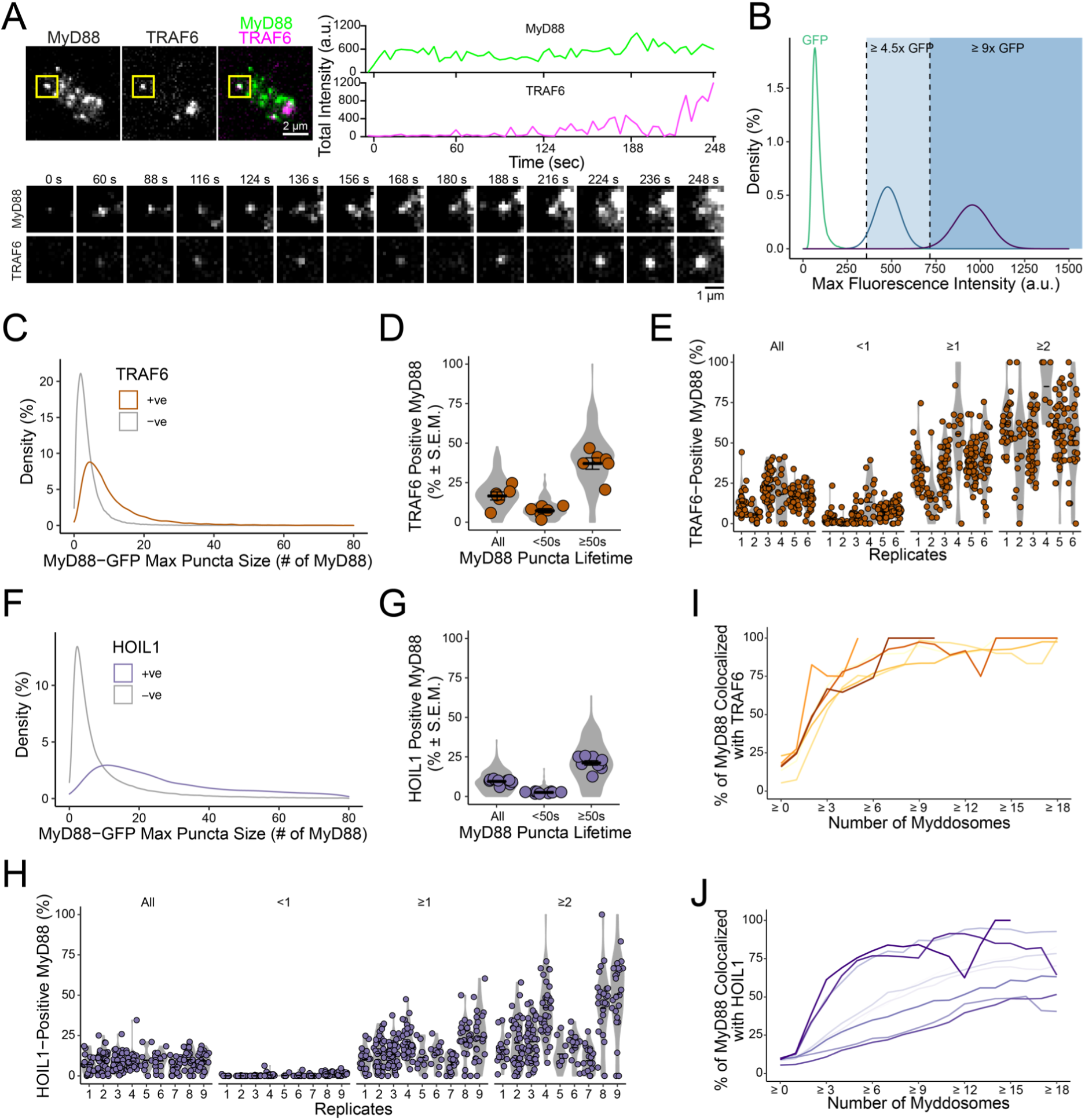
Characterization of dynamics of MyD88-GFP/mScarlet-TRAF6 and MyD88-GFP/mScarlet-HOIL1 cells. (A) TIRF images of an EL4 cell expressing MyD88-GFP and mScarlet-TRAF6. Time-series images indicate MyD88 and TRAF6 puncta from the yellow boxed area. TRAF6 is transiently recruited to MyD88 until TRAF6 becomes stable and nucleates on MyD88 puncta. Fluorescence intensities of MyD88 and TRAF6 overtime are shown at top right. (B) Density plot of single molecules of GFP (green, n = 40229 GFP particles) and estimated intensity distribution of a 6X GFP multimer (blue) and a 12X GFP multimer (purple). Shaded light blue and dark blue regions designate intensity values ≥4.5X GFP and ≥9X GFP, respectively, which were used to categorize MyD88 puncta as containing ≥1 or ≥2 Myddosome complexes. (C) Density plot showing the distribution of MyD88 oligomer size (number of MyD88-GFP monomers is derived from the maximum intensity divided by the average intensity of GFP) for MyD88 puncta that are positive (+ve) or negative (-ve) for TRAF6. The average size for puncta positive or negative for TRAF6 recruitment are 10.4 vs 3.9 MyD88s, measured from 13526 positive MyD88 puncta vs 83462 negative MyD88 puncta measured in 191 cells and combined from 6 replicates. (D) Quantification of the percentage of MyD88-GFP puncta per cell that colocalizes with TRAF6 for all puncta and puncta with lifetimes <50 s or ≥50 s. Violin plots show the distribution of individual cell measurements. Colored dots superimposed on violin plots correspond to the average value in the independent experiments (n = 6 experimental replicates, with 17-48 cells measured per replicate). Bars represent mean ± SEM. (E) Percentage (%) of MyD88-GFP puncta that colocalizes with TRAF6 per cell for all puncta, and puncta categorized as containing <1, ≥1, or ≥2 Myddosome complexes, for individual experimental replicates shown in Fig. 5C. Violin plots indicate the distribution of individual cell measurements. Colored dots superimposed on the violin plots are the averages from individual cell measurements (n = cells for replicates 1-6: n = 29,17,45,17,48 and 35). Bar represents mean. In total a minority of MyD88 puncta recruit TRAF6 (“All”, replicates 1-6: 13.5%, 5.3%, 23.0%, 18.1%, 16.7% and 15.9%). The percentage of TRAF6 recruitment was low for MyD88 puncta containing <1 Myddosome complex (“<1”, replicates 1-6: 3.3%, 1.0%, 3.3%, 13.3%, 7.3% and 8.4%), but increased with the number of complexes per puncta (“≥1”, replicates 1-6: 34.3%, 15.5%, 33.9%, 55.5%, 38.3% and 39.6%; “≥2”, replicates 1-6: 61.3%, 43.4%, 50.7%, 85.0%, 55.7% and 54.0%). (F) Density plot showing the distribution of MyD88 oligomer size (number of MyD88-GFP monomers is derived from the maximum intensity divided by the average intensity of GFP) for MyD88 puncta that are positive (+ve) or negative (-ve) for HOIL1. The average size for puncta positive or negative for HOIL1 recruitment are 47.4 vs 11.6 MyD88s, mean calculated from 10300 HOIL1 positive MyD88 puncta vs 108054 negative MyD88 puncta measured from 230 cells and combined across 9 replicates. (G) Quantification of the percentage of MyD88-GFP puncta per cell that colocalizes with HOIL1 for all puncta and puncta with lifetimes <50 s or ≥50 s. Violin plots show the distribution of individual cell measurements. Colored dots superimposed on violin plots correspond to the average value in the independent experiments (n = 9 experimental replicates, with 9-46 cells measured per replicate). Bars represent mean ± SEM. (H) Percentage (%) of MyD88-GFP puncta that colocalizes with HOIL1 per cell for all puncta, and puncta categorized as containing <1, ≥1, or ≥2 Myddosome complexes, for individual experimental replicates shown in Fig. 5C. Violin plots indicate the distribution of individual cell measurements. Colored dots superimposed on the violin plots are the averages from individual cell measurements (n = cells for replicates 1-9: n = 19, 35, 46, 34, 9, 15, 21, 27 and 24). Bar represents mean. In total a minority of MyD88 puncta recruit HOIL1 (“All”, replicates 1-6: 7.0%, 9.3%, 8.4%, 9.1%, 9.7%, 9.2%, 5.5%, 10.0% and 9.2%). The percentage of HOIL1 recruitment was low for MyD88 puncta containing <1 Myddosome complex (“<1”, replicates 1-6: 0.06%, 0.01%, 0.04%, 0.52%, 0.26%, 0.34%, 0.11%, 1.2% and 1.4%), but increased with the number of complexes per puncta (“≥1”, replicates 1-6: 10.2%, 13.8%, 13.0%, 21.1%, 11.4%, 12.8%, 6.8%, 20.9% and 24.1%; “≥2”, replicates 1-6: 16.2%, 20.4%, 21.1%, 42.2%, 13.5%, 17.4%, 9.4%, 41.9% and 50.7%). (I and J) Quantifications of percentage of TRAF6-(I) or HOIL1-(J) positive MyD88 puncta as a function of Myddosome numbers per MyD88 puncta across experimental replicates shown in Fig. 5E.

**Figure S7.**
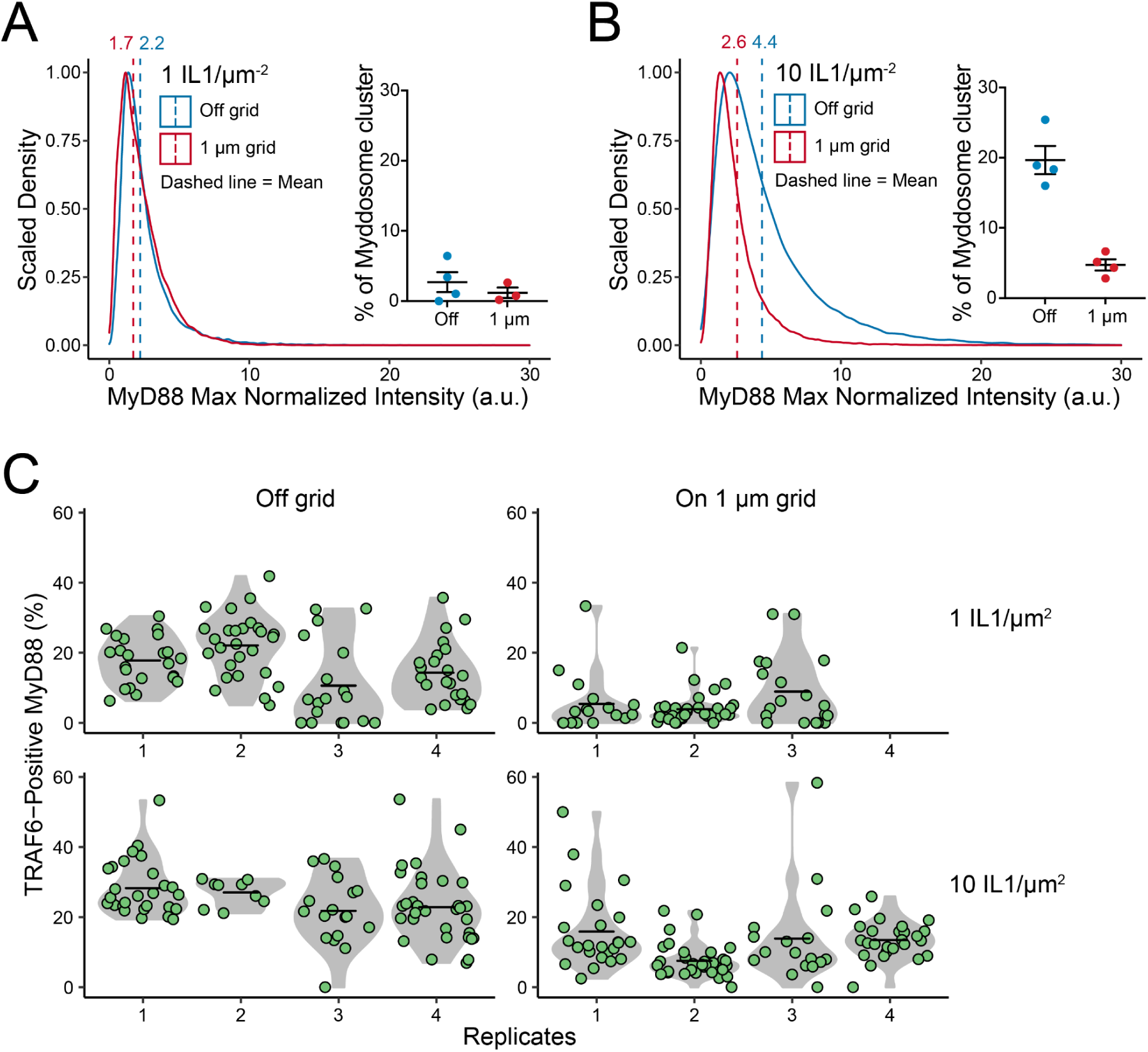
Characterization of dynamics of MyD88-GFP/mScarlet-TRAF6 cells off grids and on 1 µm grids. (A and B) Scaled density distribution of MyD88 max normalized intensity off grids and on 1 µm grids at a ligand density of 1 (A) or 10 (B) IL1/µm^2^. The average MyD88 max normalized intensity (dashed line) at 1 IL1/µm^2^ off grids versus on 1 µm grids are 2.2 versus 1.7, and at 10 IL1/µm^2^ are 4.4 versus 2.6. Insets are quantifications of the percentages of Myddosome clusters. A Myddosome cluster is defined as a MyD88-GFP puncta containing equal to or greater than 2 Myddosomes. At 1 IL-1/µm^2^, the percentages of Myddosome clusters off grids versus on 1 µm grids are 2.7 ± 1.4% versus 1.2 ± 0.7%, and at 10 IL1/µm^2^ are 19.7 ± 2.0% versus 4.7 ± 0.8%. Bars represent mean ± SEM. At 1 IL1/µm^2^, data are measured from 24315 MyD88 puncta off grids from 91 cells and 4 replicates, and 23161 MyD88 puncta on 1 µm grids from 70 cells and 3 replicates. At 10 IL1/µm^2^, data are measured from 34452 MyD88 puncta off grids from 87 cells and 4 replicates, and 71525 MyD88 puncta on 1 µm grids from 100 cells and 4 replicates. (C) Percentage of MyD88-GFP puncta that colocalizes with TRAF6 off grids and on 1 µm grids at 1 or 10 IL1/µm^2^ per cell across experimental replicates shown in Fig. 6C and 6F. Violin plots indicate the distribution of individual cell measurements. Colored dots superimposed on the violin plots are the averages from individual cell measurements. Bars represent means.

**Figure S8.**
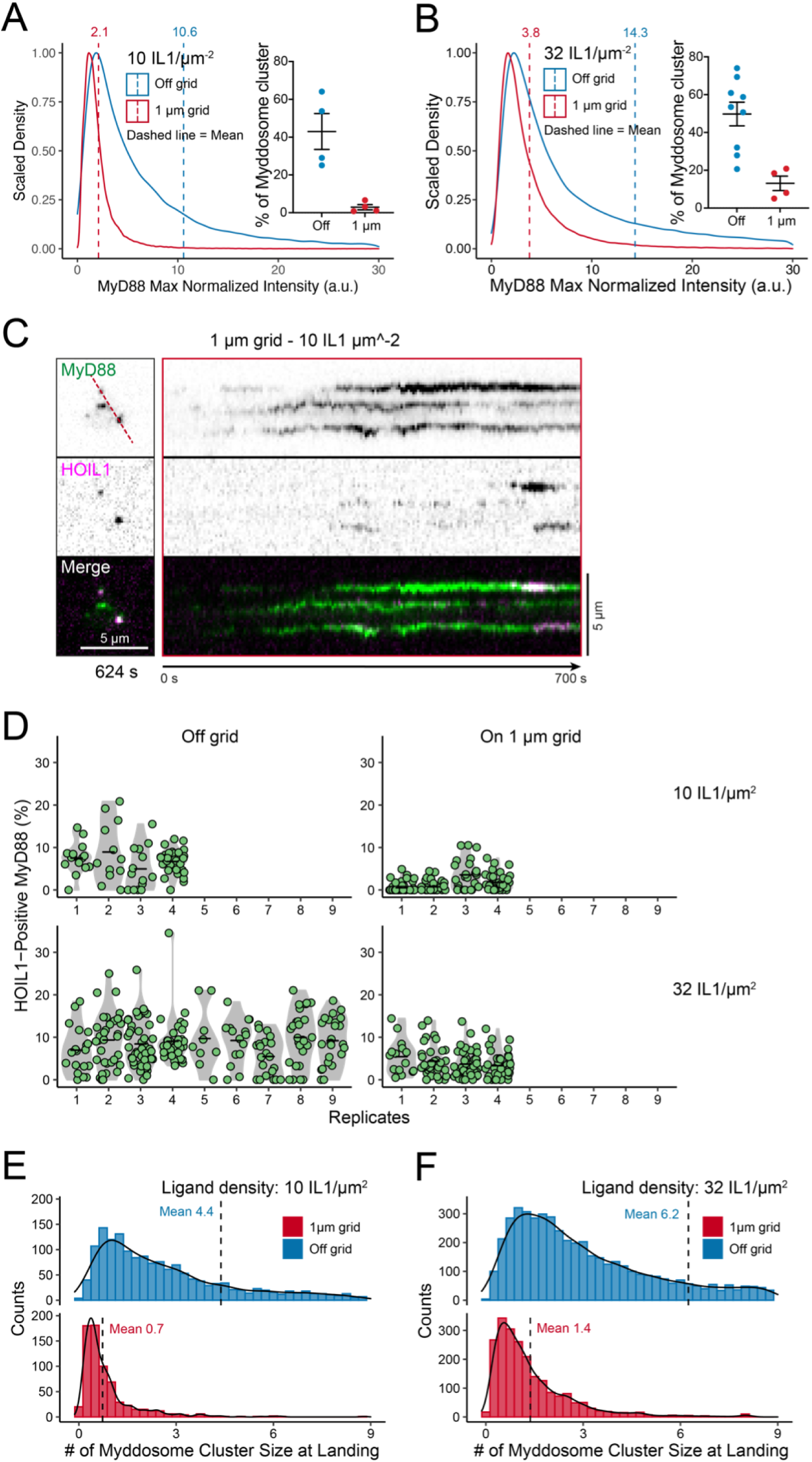
Characterization of dynamics of MyD88-GFP/mScarlet-HOIL1 cells off and on 1 µm grids. (A and B) Scaled density distribution of MyD88 max normalized intensity off grids and on 1 µm grids at a ligand density of 10 (A) or 32 (B) IL1/µm^2^. The average MyD88 max normalized intensity (dashed line) at 10 IL1/µm^2^ off grids versus on 1 µm grids are 10.6 versus 2.1, and at 32 IL1/µm^2^ are 14.3 versus 3.8. Insets are quantifications of the percentage of Myddosome clusters. A Myddosome cluster is defined as a MyD88-GFP puncta containing equal to or greater than 2 Myddosomes. At 10 IL1/µm^2^, the percentages of Myddosome clusters off grids versus on 1 µm grids are 43.0 ± 9.5% versus 2.9 ± 1.3%, and at 32 IL1/µm^2^ are 49.8 ± 6.3% versus 13.1 ± 3.9%. Bars represent mean ± SEM. At 10 IL1/µm^2^, data are measured from 53852 MyD88 puncta off grids from 74 cells and 4 replicates, and 55075 MyD88 puncta on 1 µm grids from 154 cells and 4 replicates. At 32 IL1/µm^2^, data are measured from 118354 MyD88 puncta off grids from 230 cells and 9 replicates, and 68819 MyD88 puncta on 1 µm grids from 138 cells and 4 replicates. (C) An example of TIRF images of HOIL1 recruitment in EL4 cells expressing MyD88-GFP and mScarlet-HOIL1 stimulated on IL1 functionalized SLBs on 1 µm grids at a ligand density of 10 IL1/µm^2^. Kymographs derived from dashed lines overlaid TIRF images (left panel). Scale bar, 5 µm. (D) Percentage of MyD88-GFP puncta that colocalizes with TRAF6 off grids and on 1 µm grids at 1 or 10 IL1/µm^2^ per cell across experimental replicates in Fig. 6I and 6L. Violin plots indicate the distribution of individual cell measurements. Colored dots superimposed on the violin plots are the averages from individual cell measurements. Bars represent means. (E and F) Histogram of the average landing size for HOIL1 at 10 IL1/µm^2^ (H) and 32 IL1/µm^2^ (I) off and on 1 µm grids, overlaid with density plots of the distribution. The landing size of Myddosome is calculated with landing size of MyD88 puncta divided by the intensity of 4.5xGFP. The average landing size of Myddosome at 10 IL1/µm^2^ off grids versus on 1 µm grids are 4.4 ± 1.2 versus 0.7 ± 0.2 Myddosomes (Mean ± SEM), measured from 1762 versus 691 MyD88-GFP puncta from 66 versus 86 cells and 4 versus 4 replicates. The average landing size of Myddosome at 32 IL1/µm^2^ off grids versus on 1 µm grids are 6.2 ± 1.1 versus 1.4 ± 0.2 Myddosomes (Mean ± SEM), measured from 5562 versus 2189 MyD88-GFP puncta from 212 versus 124 cells and 9 versus 4 replicates.

**Figure S9.**
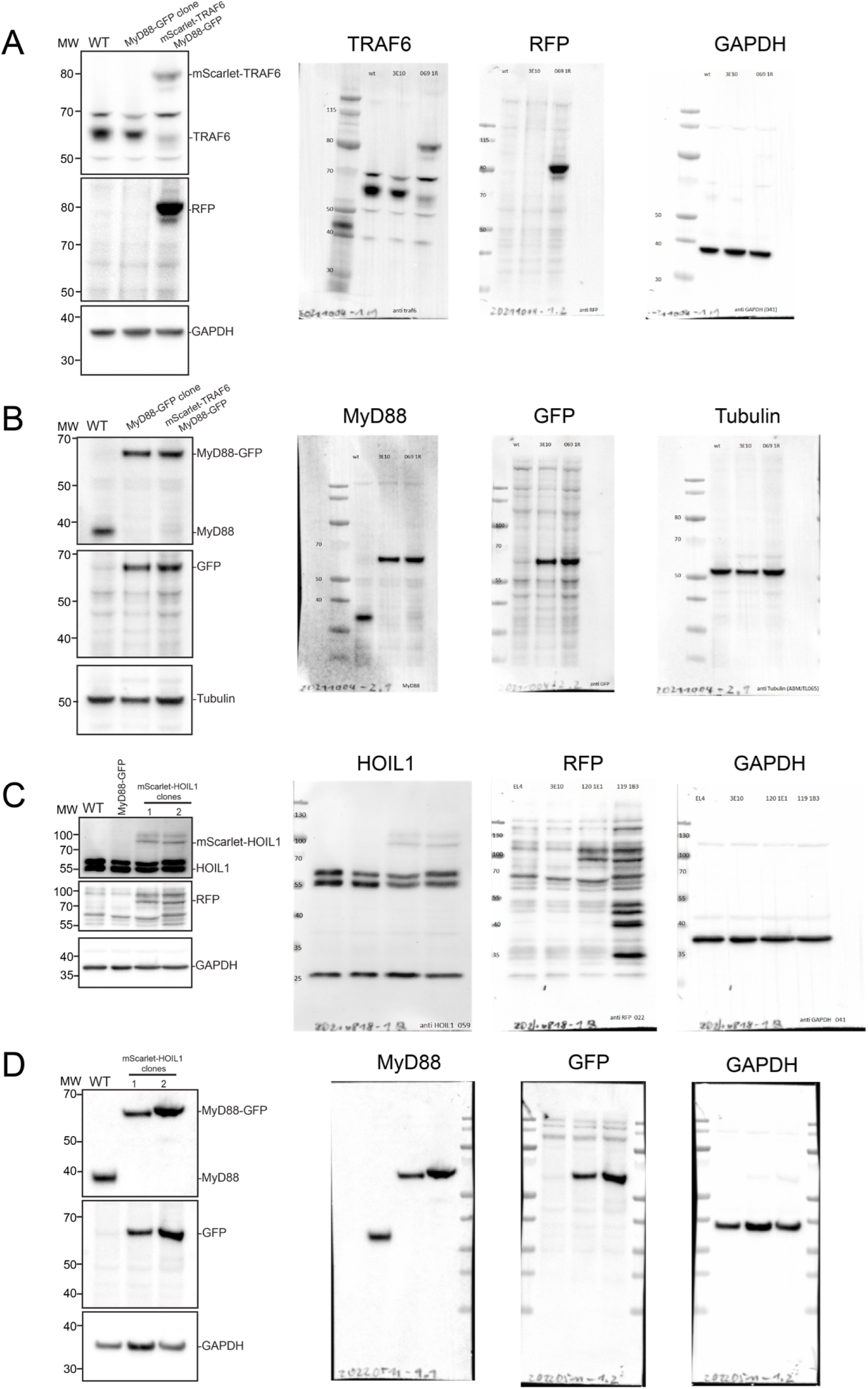
Full western blots. (A and B). Full western blots of MyD88-GFP/mScarlet-TRAF6 cell line, from figure S5A. (C and D). Full western blots of MyD88-GFP/mScarlet-HOIL1 cell line, from figure S5B.

**Table S1.**
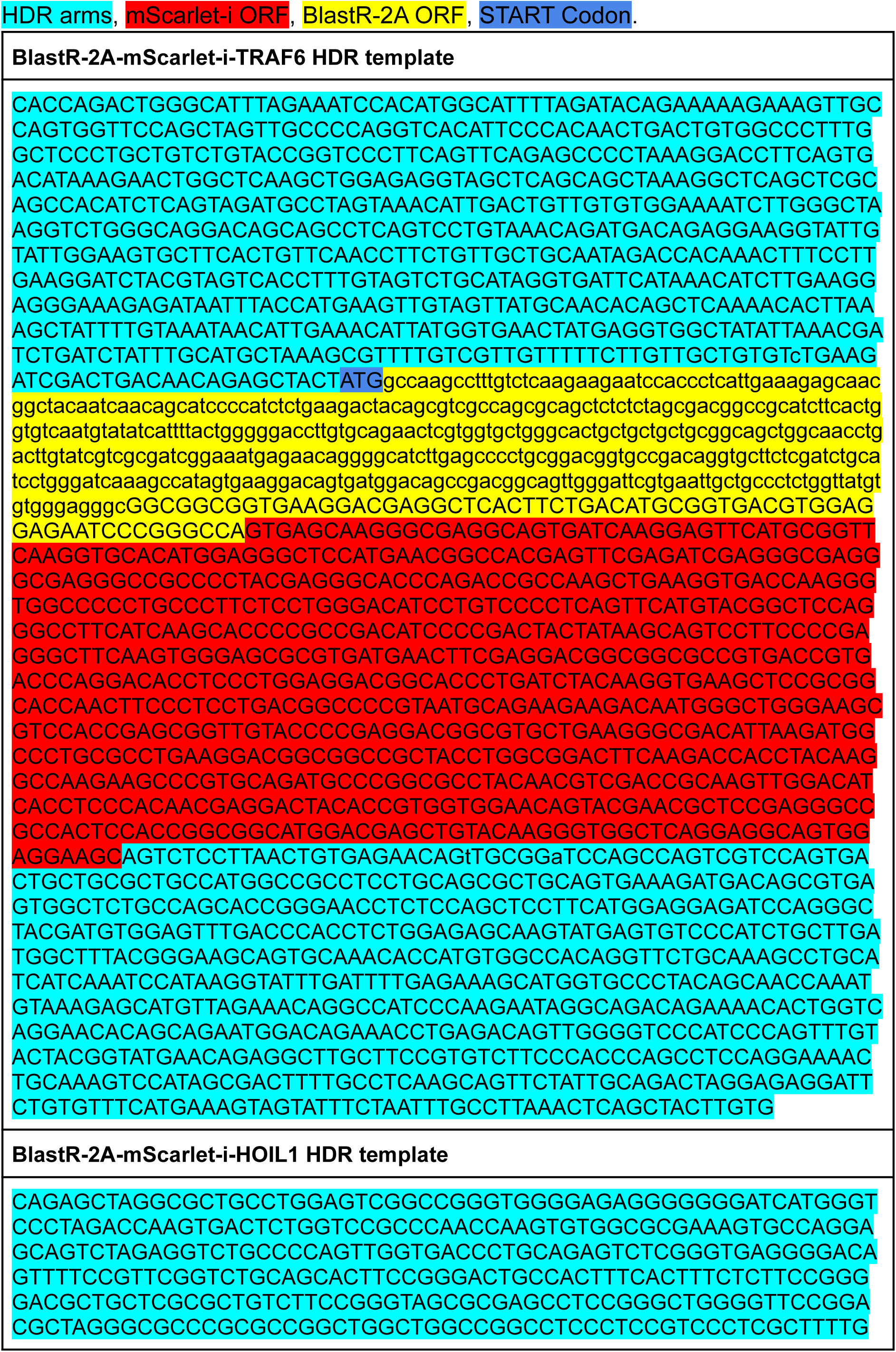
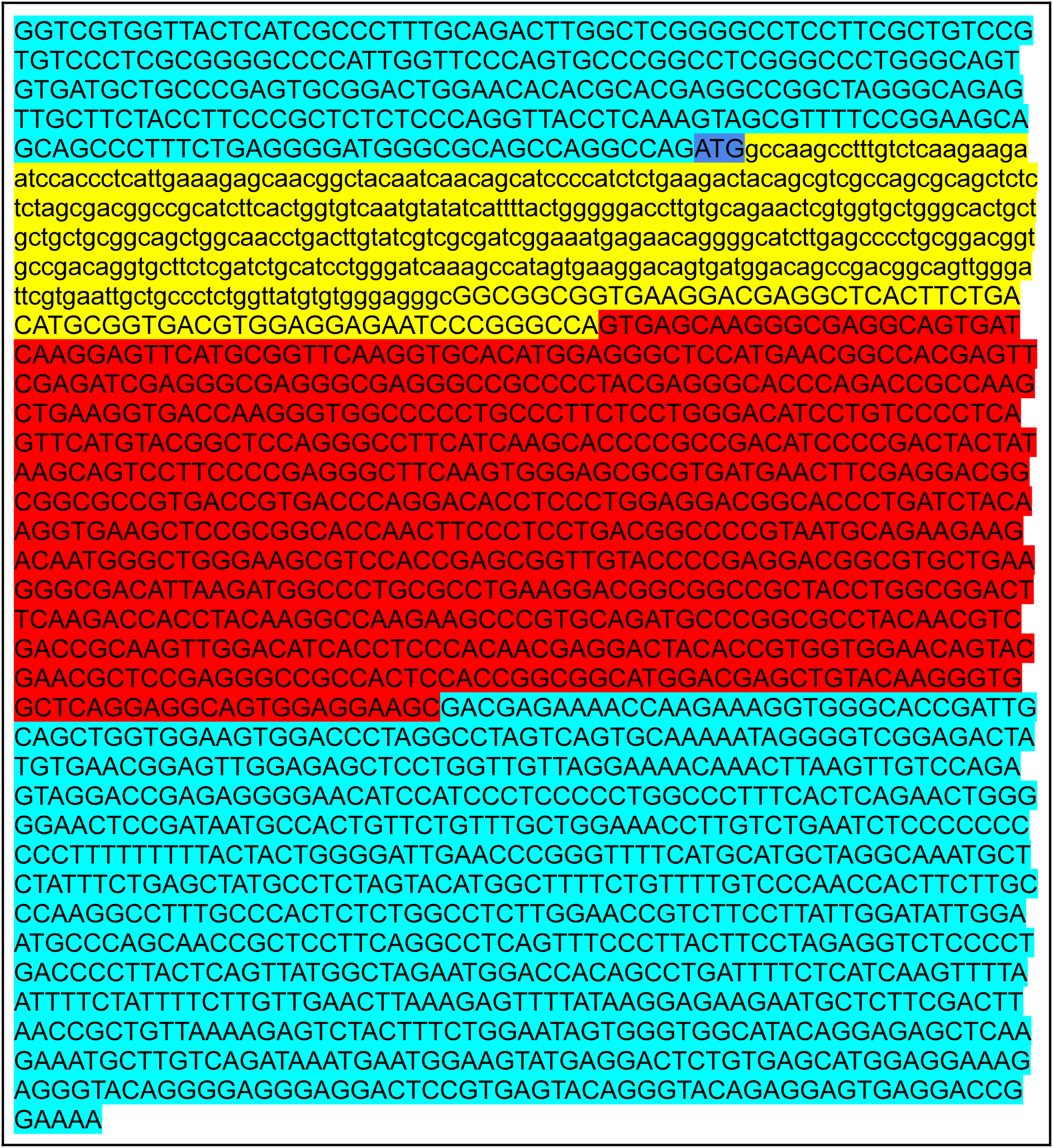
Sequences of the HDR templates

